# Tuft dendrites in frontal motor cortex enable flexible learning

**DOI:** 10.1101/2025.03.13.642781

**Authors:** Eduardo Maristany de las Casas, Kris Killmann, Moritz Drüke, Lukas Münster, Christian Ebner, Robert Sachdev, Dieter Jaeger, Matthew E. Larkum

## Abstract

Flexible learning relies on integrating sensory and contextual information to adjust behavioral output in different environments. The anterolateral motor cortex (ALM) is a frontal area critical for action selection in rodents. Here we show that inputs critical to decision-making converge on the apical tuft dendrites of L5b pyramidal neurons in ALM. We therefore investigated the role of these dendrites in a rule-switching paradigm. Activation of dendrite-inhibiting layer 1 interneurons impaired relearning, without affecting previously learned behavior. Remarkably, this inhibition profoundly suppressed calcium activity selectively in dendritic shafts but not spines while reducing burst firing. Moreover, excitatory synaptic inputs to tuft dendrites exhibited rule-dependent clustering. We conclude that active dendritic integration is a key computational component of flexible learning.

## Main Text

Flexible, adaptive behavior is a hallmark of intelligent systems, which facilitates optimal learning in dynamic, ever-changing environments. It requires continuous integration of sensory information and the ability to optimize motor plans in a context-dependent manner. In rodents, one key site of integration for these information streams is thought to be anterolateral motor cortex (ALM), a frontal cortical area highly involved in perceptual decision-making (*1*). Here, layer 5b pyramidal extratelencephalic (L5b ET) neurons encode both motor preparation and execution of choice behavior (*2*). Thus, these cells need to credit and integrate different sensorimotor combinations to adjust behavioral output in a rule-dependent manner. One common characteristic shared by all L5b ET neurons cortex-wide, including those in ALM, is having two main compartments for integrating synaptic input (basal and apical), which we hypothesize is a fundamental property of cortical architecture (*3*). According to this hypothesis, the apical compartment is endowed with powerful non-linear properties, such as NMDA- and calcium spikes, that have been proposed as plausible mechanisms for flexible learning required for adaptive behavior (*4–8*). Interestingly, the apical dendritic compartment in ALM receives inputs from sensory areas such as S1 (originating from neurons in layers 5a and 6b) and motor-related thalamic areas such as VM/VL, which target layer 1 of ALM (*9, 10*; FigS1). This input structure contrasts with apical dendritic input in layer 1 of sensory areas which tends to be top-down long-range feedback information (*11*). While layer 1 computations in sensory areas have been investigated and related to perception and learning, their role in frontal areas such as ALM remains poorly understood (*12–15*).

### Activation of layer 1 inhibition in ALM disrupts rule-dependent flexible learning

We designed a behavioral paradigm to study rule-dependent flexible learning during decision-making (Fig. 1A; adapted from *1, 16, 17*). Here, we trained animals to switch between two different decision-making tasks. First, animals were trained to follow a relatively difficult discrimination task, where the animal had to lick for a reward in response to an auditory cue arriving 1 s after a mild air-puff stimulation of the left or right whisker pad. In this task (Rule A), the animal had to lick the side corresponding to the instruction side (Fig. 1A, left). After animals became experts in this task (typically up to two weeks of training), the rule-switching paradigm was initiated, in which the animals had to switch from Rule A to the simpler Rule B, where the rewarded side was always on the left regardless of the instruction side of the air-puff (Fig. 1A, left). Finally, the animals were trained to switch back to Rule A (A’). The rule-switching paradigm occurred over 5 sessions (Fig. 1A, right). Thus, this paradigm was designed to test mechanisms involved in updating and recalling learned rules.

**Fig. 1.**
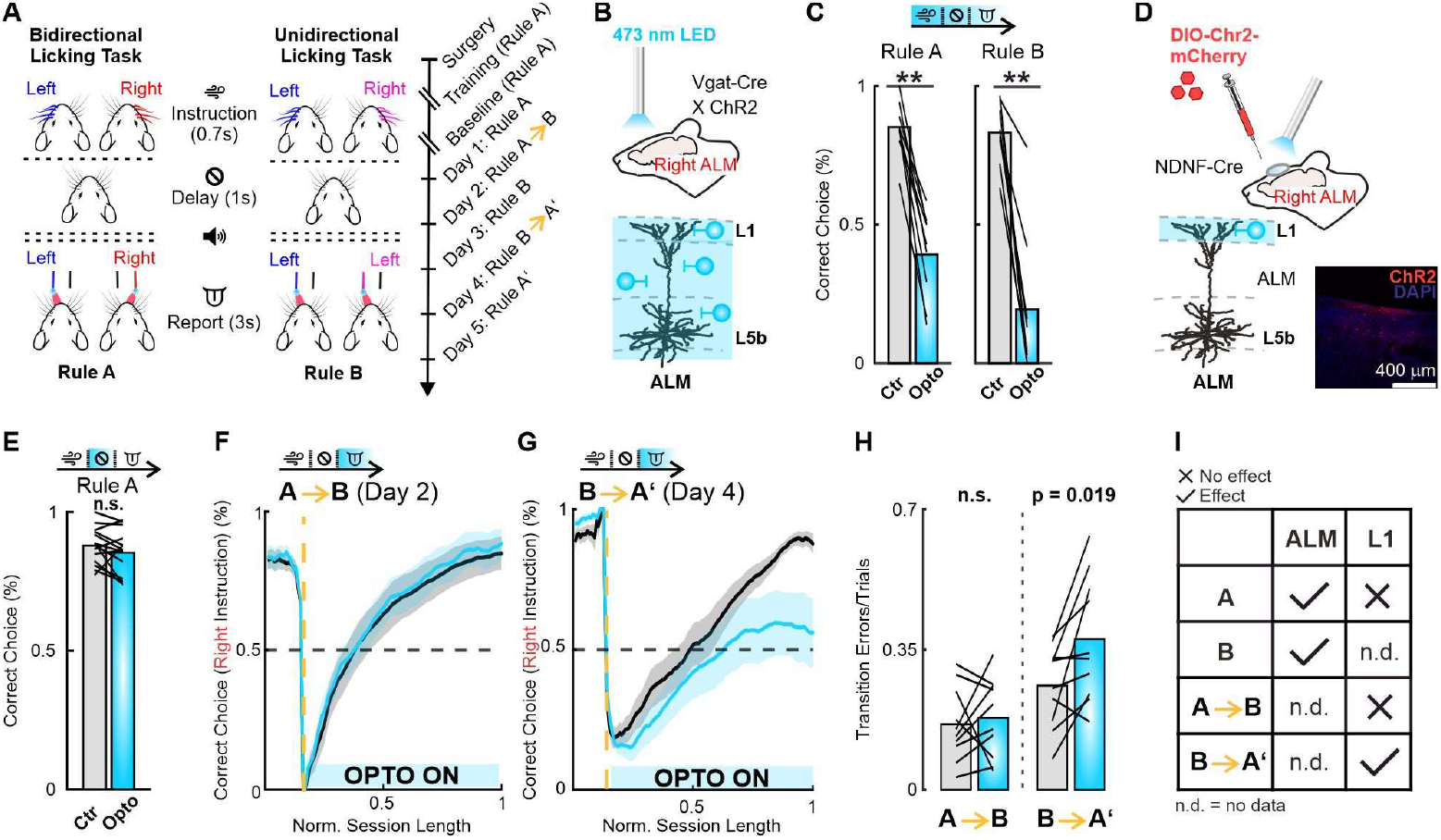
Layer 1 inhibition in ALM disrupts rule-dependent flexible learning. **(A) Left:** Behavioral task design. **Right:** Timeline of experimental procedures. **(B)** Experimental design for optogenetic silencing of ALM. Optogenetic stimulation (right hemisphere; Fig. S2 for details) through cleared skull (see Methods) to activate all interneuron types (Vgat^+^ cells). **(C)** Performance with and without ALM inhibition during all trial epochs(Rule A: Wilcoxon signed rank test, p = 0.0039, 9 sessions, 3 animals; Rule B: Wilcoxon signed rank test, p = 0.0040, 8 sessions, 3 animals). See Fig. S2 for details. **(D)** Experimental setup for optogenetic activation of NDNF interneurons during the relearning paradigm. **Inset:** Expression pattern of L1 NDNF interneurons in ALM. **(E)** Performance with and without NDNF inhibition during the delay period (one-way ANOVA, p = 0.89, 21 sessions, 10 animals). See Fig. S3 for details. **(F)** Moving average of task performance (blocks of 20 trials) throughout Day 2, transitioning from Rule A to B (right-instruction trials only). Yellow dashed line marks the introduction of the rule switch. Optogenetic (cyan) compared to control (black). ALM was activated during the report epoch (3 seconds) Mean ±s.e.m. **(G)** Same as in (F) for the Rule B to A’ switch on Day 4. Mean ±s.e.m. **(H)** Transition errors per trial during Rule A to B and Rule B to A’ switches (A-B switch: Wilcoxon signed rank test, p = 0.695, 10 mice; B-A’ switch: Wilcoxon signed rank test, p = 0.019, 10 mice). **(I)** Summary table of all inactivation experiments in Fig. 1.

Our first aim was to confirm that ALM is involved in both tasks of the rule-switching paradigm. To test this, we used a Vgat-Cre X ChR2 mouse line to optogenetically inhibit ALM for the whole trial duration (6 seconds, Fig. 1B), which impaired performance in both tasks (Fig. 1C). This effect was observed with both uni- and bilateral stimulation for both tasks (Fig. S2, see Supplement Text for detailed description).

Next, we sought to examine the contribution of the apical dendritic compartment to flexible learning by optogenetically activating dendrite-targeting inhibitory neurons, namely neuron-derived neurotrophic factor (NDNF) interneurons (Fig. 1D). The axons of these interneurons primarily innervate layer 1, activating both GABA-A and GABA-B receptors on the apical tuft dendrites of pyramidal neurons, which has been shown to reduce dendritic supralinearities like calcium spikes (*12, 13*). Activating dendritic inhibition during the delay period (decision epoch, 1 second, (*1*)) had no effect on learned behavior (rule A), per se (Fig. 1E; also see Fig. S3A&B for other epochs and more details). To test the effect of NDNF activation during rule switches (A → B; B → A’), each animal underwent the complete relearning paradigm twice; once under control conditions and once under optogenetic stimulation (random order; Control VS. Stimulated; Fig S4). During rule switches (A → B; B → A’), the animal had to change its association by switching the response side to right air-puffs (but not to left air-puffs). We therefore focused on the performance in right air-puff trials to assess the ability of the mice to switch rules. Immediately after the rule change, the performance in right-puff instructed trials dropped abruptly, but recovered to previous performance levels within a session (Fig. 1F&G; black traces). In contrast, performance in left-puff instructed trials, which did not change its meaning, remained unchanged (Fig. S3C-E). To test the effect of dendritic inhibition on relearning, we activated NDNF interneurons during the reward phase (the 3 seconds epoch in which the animal assessed trial outcome). Interestingly, although there was no effect of dendritic inhibition on switching from the more complex Rule A to the simpler Rule B, activation of NDNF interneurons drastically decelerated the animal’s relearning when shifting back from Rule B to recall the more complex Rule A’ (Fig. 1F-H). Moreover, even though optogenetic manipulation was discontinued, the effect of dendritic inhibition in the previous rule switching session persisted on the subsequent day 5 (Fig. S3F), emphasizing the manipulation’s effect on learning. In summary, while activating broad-based inhibition throughout ALM was effective in reducing performance in both rules when already learned, specific activation of NDNF inhibitory neurons profoundly affected the relearning of the more complex rule (Fig. 1I).

### Layer 1 interneuron activation prevents bursting and calcium activity in dendritic shafts

Next, we sought to understand the effect of NDNF-mediated dendritic inhibition by imaging calcium activity in apical dendrites of L5b ET cells while simultaneously exciting NDNF interneurons chemogenetically. We combined two viral strategies, one to selectively express Cre-dependent excitatory DREADDs in NDNF neurons and one to express GCaMP6s in layer 5b ET cells by injecting a retrograde Flp-dependent virus in the medulla (Fig 2A&B; see Methods). For this experiment, we performed whisker stimulation - that we expected to lead to synaptic input to layer 1 of ALM (Fig S1) - while imagining spines and dendritic shafts (Fig. 2A&B, Fig S5A). Under control conditions, dendritic shafts and spines exhibited calcium activity, both spontaneously and evoked by stimulation of the contralateral whiskerpad (Fig. 2C, D & S5B, black traces). Remarkably, NDNF-mediated inhibition induced by DCZ application, virtually eliminated calcium activity in dendritic shafts but only slightly affected spines (Fig. 2C-F and S5B, cyan traces).

**Fig. 2.**
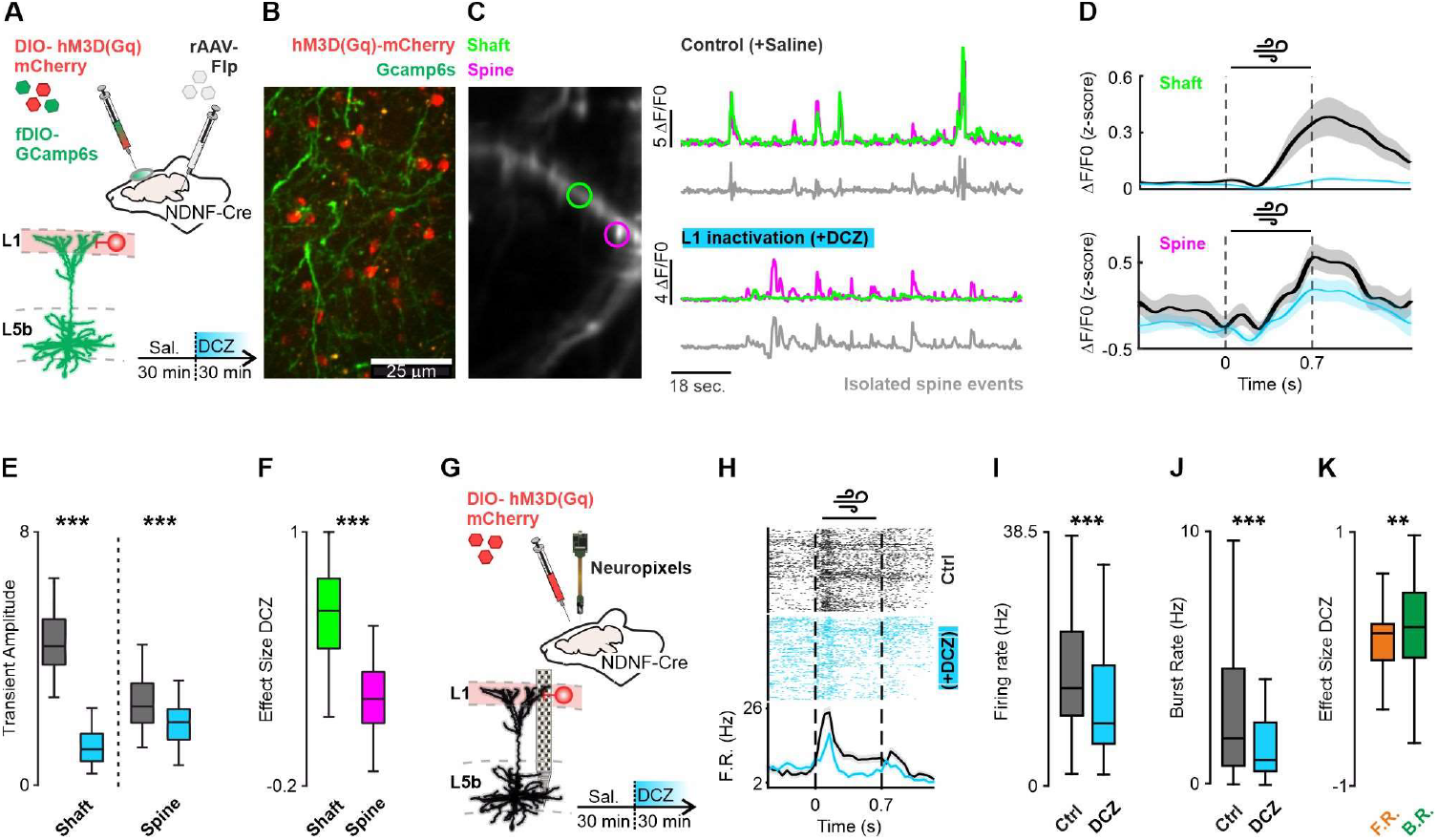
Activation of NDNF interneurons suppresses dendritic shaft calcium activity and somatic bursting. **(A)** Viral injection strategy for selective co-expression of GCaMP6s in L5b cells and excitatory DREADDs in L1 NDNF interneurons. **(B)** 2-photon averaged z-stack showing GCaMP6s expression in L5b cell dendrites and hM3D(Gq)-mCherry in NDNF interneurons in layer 1. **(C) Left:** Functional 2P imaging of L5b dendritic branches expressing GCaMP6s. **Right:** Example activity traces for shafts (green) and spines (magenta) under saline (top) or DCZ application (bottom). Grey traces show isolated spine events after subtracting shaft activity (*see Methods*). **(D) Top:** Average activity for example dendritic shaft while performing contralateral whisker stimulation. Control (black), activation of NDNF interneurons via DCZ (cyan). **Bottom:** Same for an exemplary spine. Mean±s.e.m. **(E)** Transient amplitudes for spines and shafts under saline (grey) and DCZ (cyan) (Mann Whitney U test, p < 0.0001; 52 branches, 122 spines; 3 mice). **(F)** Effect size (*see Methods*) of DCZ on calcium activity in spines and shafts (Mann Whitney U test, p < 0.0001; 3 mice). **(G)** Chemogenetic activation of NDNF interneurons during Neuropixels recording in ALM. **(H)** *Top:* Raster plot showing an illustrative firing pattern of a L5 cell during whisker stimulation. Each row is one trial; Black rows (control); cyan rows (activation of NDNF interneurons via DCZ). *Bottom:* Peri stimulus time histograms (PSTH). Mean±s.e.m. **(I)** L5 cell firing rates under control and chemogenetic conditions (Wilcoxon signed rank test, p<0.0001; 72 layer 5 neurons, 5 animals). **(J)** Same analysis for burst rates (Wilcoxon signed rank test, p<0.0001; 72 layer 5 neurons, 5 animals). **(K)** Effect size comparison of DCZ application for burst and firing rates (Wilcoxon signed rank test, p=0.002; 72 layer 5 neurons, 5 animals).

In awake mice, calcium transients in the apical dendritic shaft of pyramidal neurons have been shown to be mostly elicited by global dendritic events, which have been associated with backpropagating action potentials or somatic bursting (*18;* see also Fig. 4A-C). It was therefore important to test whether the abolition of calcium activity in dendritic shafts caused by NDNF activation was associated with the suppression of somatic firing or due to the block of voltage-gated calcium and/or NMDA channels via metabotropic GABA-B receptor activation in dendritic shafts (*19–21*). Thus, we recorded the somatic firing of layer 5 neurons in ALM using multielectrode extracellular recordings (Neuropixels probes; see Methods) before and after chemogenetic activation of NDNF interneurons (Fig. 2G). This manipulation elicited a moderate reduction in somatic firing that however could not account for the massive effect observed on calcium activity in the shafts of apical dendrites (Fig. 2H, I, Fig S5C; reduction_firing rate_=29%; reduction_Calcium activity_=77%). Thus, we conclude that, despite its effect on somatic firing, NDNF activation has an additional profound impact on calcium signaling in dendritic shafts, probably influencing crucial plasticity mechanisms involved in learning.

Depolarization of the apical dendritic tufts of pyramidal neurons leads to activation of non-linear processes that cause these cells to fire in bursts of action potentials (*3*). We therefore analyzed the effect of NDNF cell activation (DCZ application) on somatic bursting. Here, we found that DCZ application significantly decreased burst firing during whisker stimulation (Fig. 2J, Fig S5 D). Remarkably, the effect size on burst firing was significantly higher than for the firing rate of single action potentials (Fig. 2K). Taken together, these results indicate that NDNF interneuron activation mainly prevents dendritic shaft calcium activity and burst firing, presumably by activation of metabotropic GABA-B receptor in dendritic shafts (*19*).

### Excitatory layer 1 input to L5b cells shows rule- and epoch dependent functional clustering

The results so far indicate that supralinear processes in the tuft dendrites of layer 5 neurons facilitate optimal, flexible learning. We then sought to record excitatory synaptic input to individual apical dendritic branches during the rule-switching paradigm (Fig. 3A). To achieve semi-sparse expression of a glutamate activity sensor in L5b ET cells, we used a combination of diluted AAV-Flp-DIO and AAV-fDIO-iGluSNFR3-GPi injected into the right hemisphere of ALM in a Sim1-Cre mouse line (Fig. 3B). Remarkably, even single synaptic inputs displayed robust task encoding (Fig. 3C). Our analysis identified four distinct synaptic activity patterns: synapses that differentiated between trial types during (1) the instruction period (yellow), (2) the report period (choice, green), (3) both periods (white), or (4) neither (black; Fig. 3C).

**Fig. 3.**
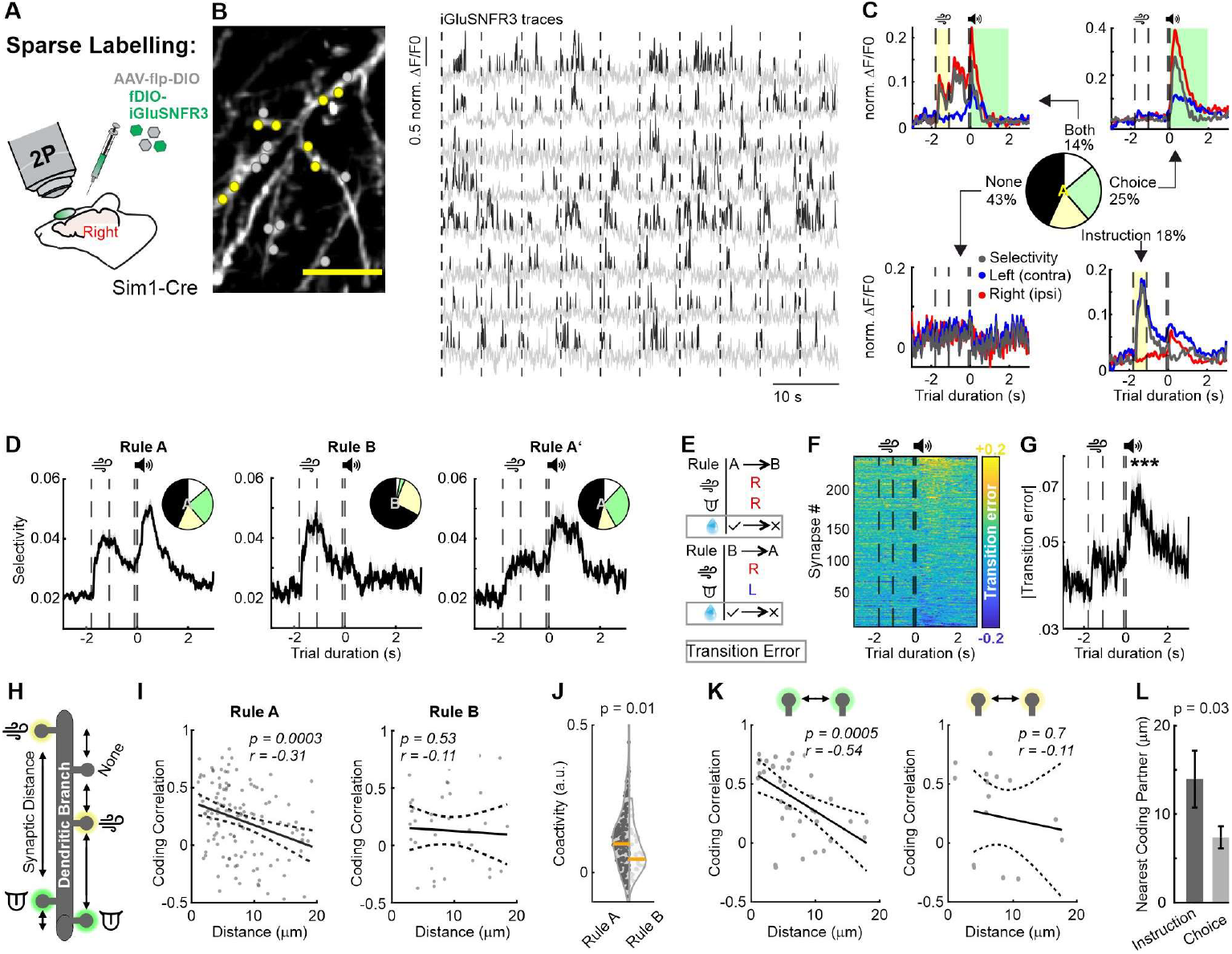
Layer 1 input to L5b ET cells shows rule- and epoch dependent functional clustering. **(A)** Viral approach combining a diluted AAV-Flp-DIO virus and a synaptic glutamate sensor (AAV-fDIO-iGluSNFR3.GPi) enabled semi-sparse labelling in L5b ET cells. **(B)** *Left*: Expression of iGluSNFR3 in apical tuft dendrites. Yellow dots indicate eight selected exemplary synapses; grey dots indicate the remaining synaptic regions of interest (ROIs). Scale bar: 30μm. *Right:* Example traces (grey) from eight synapses; putative glutamate transients (black). Dashed lines denote trial boundaries. **(C)** Average activity in left (blue) and right (red) instruction trial types from four synapses. Selectivity (grey) is defined as the absolute difference in average activity between right and left trial averages. Activity patterns are classified into significantly selective during the instruction epoch (light yellow), selective during the report epoch (light green), selective during both epochs (white), and not selective (no shading). The distribution of synaptic activity patterns is shown in the pie chart. **(D)** Average population selectivity. *Inset:* Pie chart showing the proportions of synapses with different activity patterns under different rules. Data are presented as mean ± SEM (527 synapses from all sessions, 6 animals). **(E)** Schematic visualizing the definition of transition error modulation. Synaptic activity differences for trials with the same instruction and choice conditions but differing outcomes (reward) before and after a rule switch were used to calculate transition error modulation. **(F)** Average transition error-dependent modulation during both rule switches (A→B & B→A’). Positive values indicate increased activity during transition errors, while negative values indicate decreased activity. Dashed lines denote the start of the sample period, end of the sample period, and go cue. **(G)** Average absolute transition error modulation. Data are shown as mean ± SEM (t_trial_ = - 1.8 – 0 s VS t_trial_ = 0 – 3 s; Mann–Whitney U test, p < 0.0001; 236 synapses, 6 animals). **(H)** The distance between synapses on the same branch and within 20 μm distance was measured. Synapses were classified by their activity patterns (see (C)). **(I)** Coding correlation (measure of similarity of task representation between two synapses; see Mtehods) between synapses as a function of distance. *Left:* Rule A (Spearman correlation, p = 0.0003; 131 synaptic pairs). *Right:* Rule B (Spearman correlation, p = 0.53; 38 synaptic pairs). Data are from 6 animals. **(J)** Coactivity of synaptic pairs under Rule A versus Rule B (Mann–Whitney U test, p = 0.01; 131 synaptic pairs in Rule A vs. 38 synaptic pairs in Rule B; 56 dendritic branches, 6 animals). **(K)** Coding correlation between synapses as a function of their distance. **Left:** Choice-coding synapses (Spearman correlation, p = 0.0005; 37 synaptic pairs). **Right:** Instruction-coding synapses (Spearman correlation, p = 0.70; 16 synaptic pairs). Data are from 56 dendritic branches and 6 animals. **(L)** Nearest coding partner distance for instruction- and choice-coding synaptic pairs (Mann– Whitney U test, p = 0.03; 16 instruction-selective vs. 35 choice-selective synaptic pairs, 6 animals).

To assess to what extent glutamatergic synapses discriminate between trial types, we plotted the average population selectivity (see Methods) under different rules (Fig 3D. Fig S6 A). Under rules A and A’, we found two selectivity peaks, one associated with the instruction epoch and one associated with the report epoch. In contrast, in rule B selectivity peaked only during the instruction epoch and was negligible during the report epoch (Fig 3D), although overall activity persisted throughout the trial (Fig. S6A). Thus, under rule A, a subset of excitatory inputs represented the instruction and choice side significantly, however in rule B inputs mainly represented the instruction side (Fig 3D, pie charts). This distinction aligns with the task structure, as rule A requires bidirectional licking while rule B requires only unidirectional responses (for the specific lateralization of task selectivity see Fig S9 and Supplementary Text). These findings show that task-related information is already represented in the projections of different brain areas converging in layer 1 of ALM.

During rule switches, mice learn to update their task representation when they encounter transition errors. Here, transition errors were defined as trials involving a rule change where the mouse reports the previous association and is therefore not rewarded (Fig. 3E). Thus, to determine the activity modulation caused by transition errors, we calculated the difference in synaptic activity between unrewarded, new rule and rewarded, old rule trials (Fig. 3E). We found that transition errors elicited positively and negatively regulated activity in different synaptic input populations during the report epoch (Fig. 3F). Overall, in absolute terms, synaptic input was significantly modulated by transition errors (Fig. 3G; pooled errors from A→B & B→A’), suggesting that distal tuft inputs to pyramidal neurons are important for flexible learning.

These findings suggest that crucial information from upstream areas converges in layer 1 of ALM. Functional synaptic clustering has been suggested as a mechanism to efficiently integrate different input streams (*22, 23*). We therefore examined spatiotemporal changes to synaptic input across different rules (Fig 3H). To investigate functional clustering, we determined the coding correlation (the task-dependent coactivity; see Methods) as a function of the distance between synapses sharing the same branch (Fig. 3H, I). In rule A, nearby synaptic pairs had higher coding correlation indicating functional clustering (Fig 3I). This relationship was absent in rule B (Fig. 3H, I, J; Fig S6B). In general, we found that synapses on the same branch were more coactive during Rule A than during Rule B (Fig. 3J; Fig S6B). Next, we compared synaptic clustering between choice and instruction selective synapses as their change in activity pattern during relearning differs from each other (Fig. 3D). Strikingly, we found that while coding correlation in choice selective (green label) synapses decreased as a function of synaptic distance, this relationship was not observed in instruction-selective (yellow label) synapses (Fig. 3K). Moreover, neighboring choice-selective synapses were closer together than instruction-selective synapses (Fig. 3L). In summary, we find that excitatory layer 1 input to L5b cells shows rule- and epoch dependent functional clustering. This finding suggests that there are two types of synapses at play, putatively thalamocortical (motor thalamus → ALM; presumably choice selective) and corticocortical (S1 → ALM; presumably instruction selective) which show different functional clustering and activity patterns with changing rules (see also Fig S1).

### Postsynaptic dendritic activity flexibly represents choice between rule switches

The data so far show that synaptic clustering is associated with the more complex Rule A and that dendritic inhibition, which abolished calcium transients in the shafts, also prevents optimal re-learning of Rule A. This suggests that postsynaptic mechanism(s) involved in dendritic integration underlie flexible learning (*24*). We therefore performed in vivo calcium imaging of apical dendrites with different imaging strategies to detect several forms of dendritic calcium transients. We classified dendritic events into four categories, each of them representing putative different forms of postsynaptic mechanisms: Global, Hemi-tree, branch, or spine-specific calcium events (Fig S7A). First, we used two-photon, two-plane imaging which allowed us to simultaneously track dendritic tufts and trunks and thus to detect global, hemi-tree or branch specific but not spine-specific activity (Fig. 4A, FI S7A). We imaged the same field of view (FOV) across sessions and assigned each dendritic region of interest (ROI) to its corresponding tree (Fig. 4B; tufts and trunks).

**Fig. 4.**
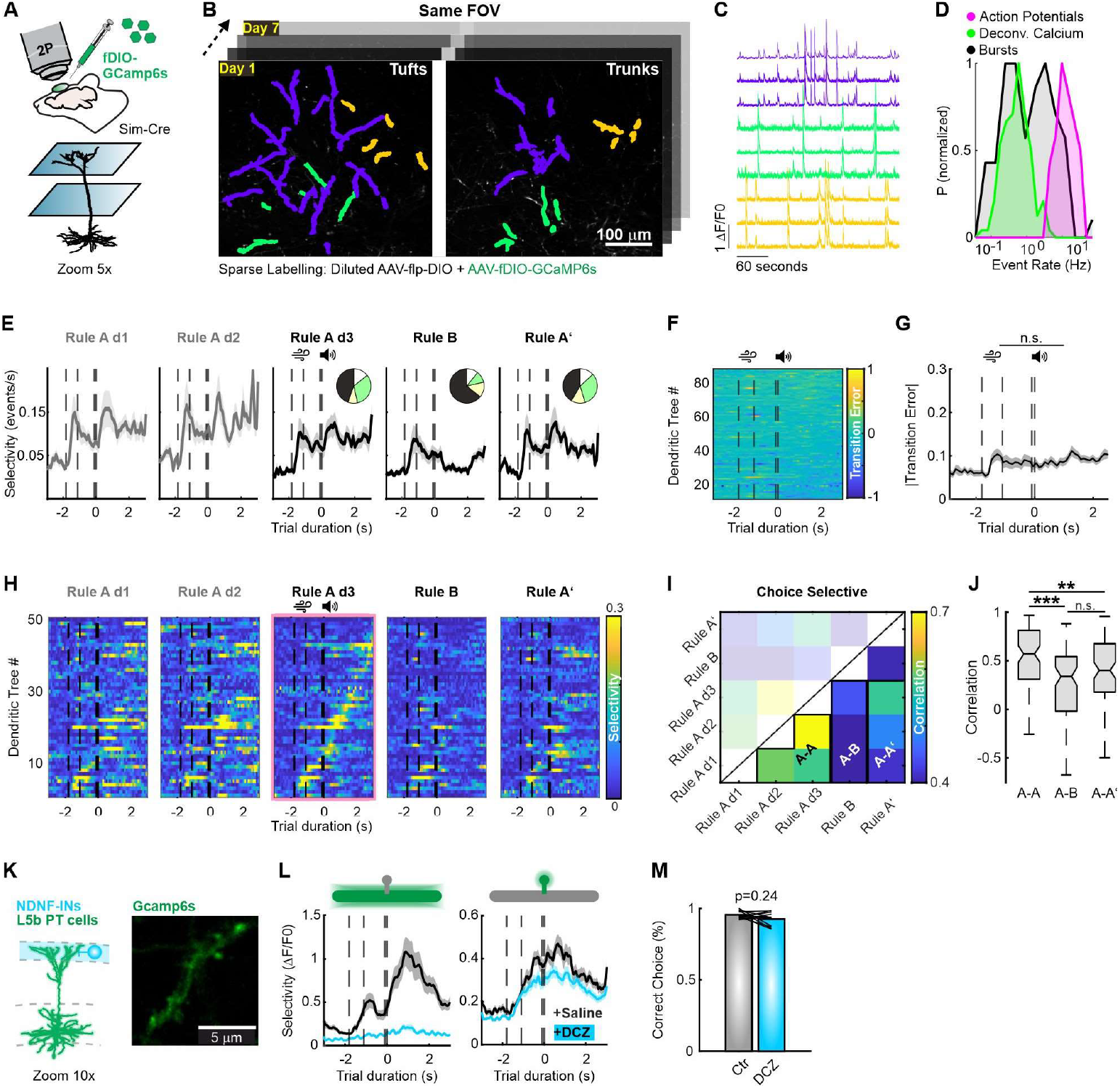
Postsynaptic dendritic activity flexibly represents choice between rule switches. **(A)** A viral approach combining diluted AAV-Flp-DIO and calcium sensor (AAV-fDIO-GCaMP6s) enabled sparse labeling in PT cells. Two-plane imaging (60 μm apart in the z-plane) was performed to capture dendritic tufts and trunks *in vivo*. **(B)** 2P imaging of the same field of view (FOV) across multiple days. Colored regions highlight example regions of interest (ROIs). Different colors represent three distinct dendritic trees from individual cells. **(C)** Calcium activity traces for three example dendritic branches (individual ROIs) from each dendritic tree. **(D)** Histogram showing the normalized event rate probability for single action potential firing (magenta), burst events (black) or deconvolved spikes derived from calcium imaging data (green). **(E)** Dendritic tree selectivity across sessions of the relearning paradigm (mean ± SEM; 51 dendritic trees, 5 animals). *Insets:* Pie charts showing the distribution of different activity patterns of dendrites across sessions. Activity patterns are classified as follows: selective during the instruction epoch (light yellow), selective during the report epoch (light green), selective during both instruction and report epochs (white), and not selective (black). Rule A: 45% not selective, 9% instruction selective, 32% choice selective, 14% both; Rule B: 64% not selective, 14% instruction selective, 11% choice selective, 11% both; Rule A’: 42% not selective, 13% instruction selective, 32% choice selective, 13% both **(F)** Transition error-dependent activity for all dendritic trees imaged during both rule switches (A→B & B→A’). Positive values indicate increased activity during transition errors, while negative values indicate decreased activity. **(G)** Average absolute transition error modulation for all dendritic trees. Data are presented as mean ± SEM (t_trial_ = -1.8 – 0 s VS t_trial_ = 0 – 3 s; Mann–Whitney U test, p = 0.07; 108 neurons, 10 sessions [5 A-B, 5 B-A’ switches], 5 animals). **(H)** Trial-type selectivity for all dendritic trees (rows) during the relearning paradigm. Trees from all sessions were sorted based on their peak selectivity timepoint at the day before the first rule switch (Rule A, day 3; third column, pink frame). **(I)** Average day-to-day coding correlation (see Methods) for single dendritic trees across relearning. Black frames denote comparisons between Rule A-A, Rule A-B, and Rule A-A’ sessions. Only choice-representing dendritic trees were analyzed. **(J)** Average coding correlation between Rule A-A, A-B, and A-A’ sessions in choice-representing dendritic trees (Friedman’s Test, p = 0.0004; Mann Whitney U test with Bonferroni correction, A-A vs. A-B, p < 0.0001; A-A vs. A-A’, p = 0.018; A-B vs. A-A’, p = 0.06; 29 choice-selective dendritic trees, 5 animals). **(K)** *Left:* Illustration of the experimental setup (referenced in Fig. 1C). *Right:* Exemplary FOV for high magnification dendritic calcium imaging. **(L)** Average selectivity of shaft events (**left**) and isolated spine events (**right**) under control conditions (saline application, black) or during NDNF interneuron activation (DCZ application, cyan). Data are shown as mean ± SEM (52 branches, 122 spines, 3 mice). **(M)** Behavioral performance comparing control versus DCZ application (Wilcoxon signed rank test, p=0.24; 11 Rule A sessions, 3 animals).

In this first dataset dendritic activity was dominated by global events that were synchronized across the entire dendritic tree (Fig. 4C), consistent with previous reports linking such events to backpropagating action potentials and/or bursting (*18, 25*; but see also *5, 26*). However, based on the comparison of our calcium and electrophysiological data, (Fig 4D) and on studies that previously have combined single cell electrophysiology with calcium imaging in vivo (18, *27*), we conclude that calcium transients recorded in apical dendrites are caused by fast trains of several action potentials i.e. putative bursts and not by single action potentials (Fig 4D). To further understand the role of these dendritic events during adaptive behavior we plotted their trial-type discrimination (selectivity) throughout the paradigm. We found that dendrites robustly exhibited high selectivity during both the instruction and report period in Rule A and A’ sessions (Fig 4E). However, once again as described for layer 1 input, selectivity was abolished under the simpler Rule B during the report period (Fig. 4E, Fig. S7B, C, for the specific lateralization of task selectivity see Fig S9 and Supplementary Text).

We next examined whether L5b dendrites were modulated by transition errors. Unlike excitatory inputs (Fig 3), dendritic activity showed no significant error modulation (Fig. 4F, G; pooled errors from A→B & B→A’), suggesting that L5b ET neurons receive synaptic information about errors related to changing associations but do not represent it in their dendritic calcium activity.

Representational drift has been suggested as a plausible mechanism to balance stability and plasticity of neural networks while avoiding interference between learned associations (*28*). To assess representational drift within and across tasks, we calculated coding correlation, a measure of task-related activity similarity (*See Methods*), for individual dendritic trees throughout the paradigm (Fig. 4H, I). Choice-selective dendrites exhibited stable coding correlation across rule A sessions, indicating a robust representation of learned behavior (Fig 4I, J). However, coding correlation significantly decreased when comparing rule A to rule B sessions, suggesting representational drift in response to task structure changes (Fig 4I, J). Notably, once animals transitioned back to rule A’, dendritic representations did not revert to their original state, as coding correlation between rule A and A’ sessions remained low (Fig 4I, J). This indicates that individual choice-selective dendrites do not fully recover their initial task representation after relearning. Moreover, instruction-selective dendrites showed no task-dependent representational drifts (Fig. S7 D, E).

Finally, to compare calcium transients in spines versus dendritic shafts we conducted high-magnification calcium imaging (Fig. 4K, Fig. S7A). These experiments revealed high frequency in spine events that were isolated by implementing an algorithm to detect local activity (Fig S5 B; *25*). Spine specific calcium transients, presumably elicited by NMDA channels, showed trial type dependent selectivity, which ramped up during the instruction period and peaked shortly after the go cue (Fig 4L, right, black trace). We observed a similar pattern in shafts transients (Fig 4L, left, black trace). Combining this imaging approach with chemogenetic NDNF interneuron activation during behavior, abolished trial type selectivity in dendritic shafts, which on the contrary was preserved in dendritic spines (Fig. 4K, L, cyan traces). Similarly to optogenetic activation (Fig 1), chemogenetic activation of NDNF interneurons in animals that already learned rule A had no significant effect on task performance (Fig. 4M). These findings describe the intricate interplay between different levels of dendritic integration and the effect of its inhibition by NDNF interneurons during behavior.

### Layer 1 NDNF interneurons reduce their activity during rule switching

Activating L1 NDNF interneurons had a dramatic effect on apical dendritic calcium, raising the question as to how NDNF neurons shape apical dendritic activity under control conditions. To examine this, we expressed the calcium sensor Gcamp6f in an NDNF-Cre-dependent mouse line which enabled us to track NDNF interneuron activity during behavior (Fig 5A). For each animal we performed longitudinal imaging of the same set of NDNF interneurons in L1 throughout the entire rule switching paradigm. Recorded calcium signals (black traces) were deconvolved (Cascade algorithm; grey traces) to estimate action potential activity (Fig. 5B). We found that NDNF interneurons in ALM showed a wide range of task-related activity patterns (Fig. 5C). Across the population of NDNF interneurons, their activity showed similar task representation to excitatory synaptic inputs (Fig 3) and postsynaptic dendritic activity (Fig 4). In short, (1) a subset of NDNF interneurons were active for both rules A & B (Fig. S8A); (2) in rule A (day1-day3) and A’, interneurons discriminated between trial types in both the instruction and report epoch (Fig. 5D); (3) in rule B, selectivity only peaked during the instruction epoch but not during the response epoch (Fig. 5D; (for the specific lateralization of task selectivity see S9 and Supplementary Text).

**Fig. 5.**
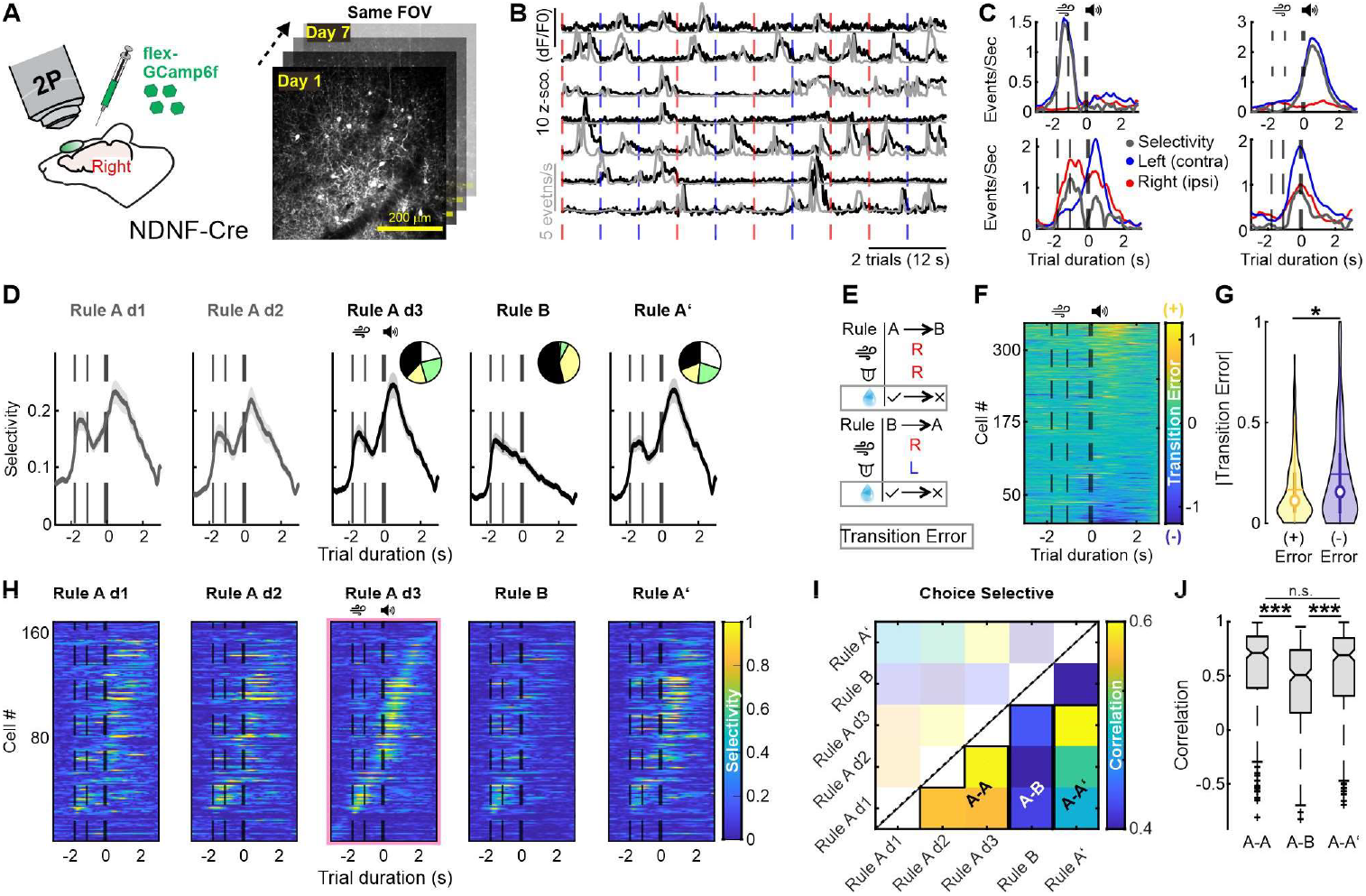
Layer 1 NDNF interneurons encode flexible decision-making and reduce activity during rule relearning. **(A) Left:** AAV-flex-GCaMP6f was injected into an NDNF-Cre mouse line to label Layer 1 (L1) NDNF interneurons. **Right:** Representative field of view (FOV). The same FOV and neurons were imaged across the entire relearning paradigm. **(B)** Example calcium activity from seven NDNF interneurons. ΔF/F0 activity (black); event-deconvolved activity (grey). Vertical dashed lines mark the onset of right (red) or left (blue) instructed trials. **(C)** Average calcium traces for left (blue) and right (red) instruction trials from four neurons. Selectivity is shown in grey. Dotted lines indicate the start of the sample period, end of the sample period, and the go cue. **(D)** Selectivity across sessions during relearning. Mean ± SEM. Insets: Pie charts illustrating the distribution of different activity patterns of neurons under various rules. Activity patterns are classified as selective during the instruction epoch (light yellow), selective during the report epoch (light green), selective during both instruction and report epochs (white), and not selective (black). Rule A: 38% not selective, 17% instruction selective, 24% choice selective, 21% both; Rule B: 54% not selective, 38% instruction selective, 6% choice selective, 2% both; Rule A’: 31% not selective, 17% instruction selective, 22% choice selective, 30% both (170 neurons from 4 animals). **(E)** Definition of transition error modulation (*see Methods*). **(F)** Averaged transition error modulation for all interneurons imaged during both rule switches (A→B & B→A’). Positive values indicate increased activity during transition errors, while negative values indicate decreased activity. Dotted lines mark the start of the sample period, end of the sample period, and the go cue. **(G)** Magnitude of transition error modulation of negatively (blue) and positively (yellow) modulated interneurons (Mann–Whitney U test, p = 0.041; 187 positively modulated neurons vs. 160 negatively modulated neurons; 12 sessions, 6 A-B and 6 B-A switches, from 4 animals). **(H)** Trial-type selectivity across the relearning paradigm. Neurons from all sessions were aligned according to their peak selectivity timepoint the day before the first rule switch (Rule A, day 3; third column, pink frame). **(I)** Day-to-day comparison of coding correlation (see Methods) for the same neurons during relearning. Black frames highlight comparisons between Rule A-A, Rule A-B, and Rule A-A’ sessions. **(J)** Average coding correlation across Rule A-A, A-B, and A-A’ sessions (Friedman’s test, p < 0.0001; all paired comparisons performed using Mann–Whitney U tests with Bonferroni corrections; 85 choice-selective neurons, 30 sessions, 4 animals).

Having established this, we went on to characterize NDNF interneurons’ representation of transition errors in rule switching sessions (Fig 5E). We found a subpopulation of NDNF interneurons being modulated by transition errors (Fig 5F; pooled errors from A→B & B→A’). Overall, NDNF interneuron activity was negatively modulated by transition errors (Fig. 5G & Fig. S8 B-D). Thus, activity of these interneurons is dampened during rule switches, potentially providing a gating mechanism for increasing dendritic supralinearities during flexible learning.

Finally, we wanted to investigate representational drift in NDNF interneurons within and between different rules (Fig. 5H; Fig. S8E). Representational drift was given by the coding correlation of each cell from session to session throughout the behavioral paradigm. In choice-selective interneurons we found high coding correlation levels within rule A before relearning (Fig. 5Hd1-d3 & I). However, coding correlation dropped significantly when comparing rule A (d1-d3) with B sessions, indicating a drift in task representation (Fig. 5I&J). Lastly, once animals switched back to A’, the task representation of choice-selective NDNF interneurons drifted back, showing high coding correlation values when comparing sessions in rule A’ and A (Fig. 5I&J). Here, unlike the drift properties of postsynaptic dendritic activity, choice selective NDNF interneurons recovered to their task representation after relearning rule A’. This task-dependent drift in representation was not observed in instruction selective interneurons, where coding correlation remained similar throughout the relearning paradigm (Fig. S8 F-G).

## Discussion

In this paper, we found evidence for active dendritic integration as a potential mechanism enabling the brain to solve the cognitive challenges posed by adaptive behavior (*24, 29*). By activating NDNF interneurons, we found that dendritic mechanisms in ALM are only necessary for the relearning of complex rules (but not for either relearning of simple rules or for the performance of an already learned task per se; Fig. 1). This implies that non-linear dendritic processes are crucial for the association of sensory with motor information, a requirement that was necessary for the more complex rule, Rule A, but not Rule B.

NDNF interneurons target extrasynaptic, metabotropic GABA_B_ receptors that have been shown to inhibit L-type calcium channels and bursting *in vitro* (*12, 19–21*). In our *in vivo* experiments, the impact of NDNF neurons on dendritic calcium was profound and has not been reported previously: the selective block of calcium signaling in shafts versus spines of L5b ET cells (Fig. 2). This effect was accompanied by a modest reduction in cell firing emphasizing its dendritic nature, and a relatively larger reduction in burst firing. We propose that flexible learning depends on the elevation of shaft calcium (via a Hebbian plasticity mechanism involving burst firing) which triggers a long-lasting enhancement of synapse-to-synapse cooperativity. This finding may also shed light on the long-standing issue of the interpretation of coherent calcium signals throughout the dendritic tree (*30*). It implies that single action potentials rarely cause calcium in apical tuft dendrites of layer 5 cells (*30*), but rather that calcium signals reflect burst firing that itself is suppressed by dendritic inhibition (Fig. 4). The data are also consistent with backpropagation-activated calcium spike (BAC) firing that involves cooperative activity in both compartments simultaneously (*30, 31*).

Strikingly, glutamate imaging revealed that task-related information is already widely represented in the upstream inputs to layer 1 of ALM (Fig. 3). Retrograde tracing showed that this activity pattern is composed of input from S1, higher-order thalamic regions and via layer 6b neurons distributed over large areas of sensory cortex (Fig. S1; *9, 10*). According to the 2-compartment view of pyramidal neurons that apical dendritic input serves as contextual information (*3*), this would imply that in ALM, input from S1 provides context for subsequent motor plans.

Functional clustering has been observed before during learning (*23, 32*). Here, we found that only synapses conveying choice, but not sensory, information were functionally clustered. We hypothesize that the two types of synapses, thalamocortical (VM → ALM; presumably choice selective) and corticocortical (S1 → ALM; presumably instruction selective), undergo different functional clustering and learning-related activity (Fig 3). A similar dichotomy in the structural composition of thalamocortical and corticocortical inputs has been observed in motor cortex over learning (*33*).

Moreover, our data point towards NDNF interneurons as putative gatekeepers for flexible learning as their activity was negatively modulated by transition errors (Fig. 5; *34, 35*). This opens the question as to what mechanisms underlie the suppression of NDNF neurons during learning? We propose increased activity of cholinergic input to cortex, which has been implied in learning paradigms (*36*). Furthermore, acetylcholine has been shown to inhibit active L1 NDNF cells by binding to muscarinic receptors (*37*) and activate VIP neurons that putatively inhibit NDNF neurons (*38*). Lastly, cholinergic input to cortical pyramidal neurons has been shown to facilitate the generation dendritic plateau potentials *in vitro* (*39*).

In conclusion, our findings highlight the importance of dendritic non-linear properties for the integration and association of different input modalities and suggests that these events are particularly important for more difficult cognitive tasks. The relearning-related changes in functional synaptic clustering point to an interplay between the active properties of shaft dendrites and the structural arrangement of excitatory inputs. Lastly, we conclude that NDNF neurons in layer 1 act as the gatekeepers for the necessary plasticity-related processes for learning.

## Acknowledgments

The authors thank Johannes Letzkus and Claudio Elgueta for helpful discussions; the members of the Larkum lab, in special those responsible for organizational tasks, for supporting this project. We thank SciDraw for multiple schematics used in this paper. We thank the colleagues of the Research Workshop at the Charité - Universitätsmedizin Berlin for developing and manufacturing the experimental devices.

## Funding

National Institutes of Health grant BRAIN Initiative 1R01 NS111470 (DJ,MEL)

European Union Horizon 2020 Research and Innovation Programme (SGA1-3: 72070/HBP, 785907/HBP, 945539/HBP, 670118/ERC ActiveCortex, 101055340/ERC Cortical Coupling to M.E.L.)

Einstein Foundation Berlin (EVF-2017-363, EVF-2017-363-2, EVF-2020-571)

Deutsche Forschungsgemeinschaft (Exc 257 NeuroCure, Grant No. LA 3442/3-1, Grant No. LA, Project number 327654276 SFB1315, Grant No. LA 3442/6-1)

## Author contributions

Conceptualization: EMC, MEL, DJ, RS

Methodology: EMC, DJ, MEL, RS, MD

Investigation: EMC, KK, MD, LM, CE

Visualization: EMC, MD

Funding acquisition: DJ, MEL, RS

Project administration: MEL, DJ, RS

Supervision: MEL, DJ

Writing – original draft: EMC, MEL, DJ

Writing – review & editing: EMC, MEL

## Data and materials availability

All data needed to evaluate the conclusions in the paper are present in the paper and/or Supplemental Materials. Source data and analysis code used in this paper will be uploaded to a public depository prior to publication.

## Supplementary Materials

Materials and Methods

Figs. S1 to S9

Supplementary Text

Table S1

## Supplementary Materials

## Materials and Methods

### Animals

All procedures are in accordance with protocols approved by the Charité – Universitätsmedizin Berlin and the Berlin Landesamt für Gesundheit und Soziales (LAGeSo) for the care and use of laboratory animals.

We used Sim1-Cre (Tg(Sim1-cre) (*40*) mice to label L5b extratelencephalic pyramidal neurons and Ndnf-tm1.1(cre)Rudy/J mice to label subpopulations of neuron-derived neurotrophic factor (NDNF) interneurons. We crossed Ai32 (129S-Gt(ROSA)26Sortm32(CAG-COP4^*^H134R/EYFP)Hze/J) transgenic mice with Vgat-Cre mice to express channelrhodopsin in all interneurons. Moreover, wild-type C57BL/6J mice were used.

### Surgical procedures

Mice were anaesthetized with isoflurane (1.5–2.0% in O_2_) or ketamine (100 mg kg−1)/xylazine (10 mg kg−1) and kept on a thermal blanket.

For retrograde fluorescent labeling, 1% Fast Blue (Polysciences) in saline was applied to a craniotomy (∼0.5 mm in diameter) made over the right ALM cortex of wild-type C57BL/6J mice. After 7 min of incubation, the surface was rinsed with saline. Mice were housed for a week before being used for histology.

For *in vivo* two-photon Ca^2+^ imaging, glutamate imaging and optogenetic manipulation we combined implantation of cranial windows with viral injections. First, the scalp and periosteum were carefully removed. A 3.5 mm craniotomy was made over ALM cortex of the right hemisphere using a biopsy punch (Stiefel). Next, viral injections were performed slowly injecting viral solution at a speed of 20nl/min. Coordinates to target ALM were +2.5 mm anterior-posterior (AP) and 1.5 mm mediolateral (ML). Depth varied based on the target cell type. For L5b ET cells virus was injected at a depth of 0.7 mm dorsoventral (DV), for targeting NDNF interneurons at a depth of 0.1 mm. For two-photon Ca^2+^ imaging of sparsely labelled L5b ET cells we injected 70 nl of AAV-Ef1a-fDIO-GCaMP6s (1×10^12^ vg/mL, Addgene) combined with AAV9-EpEF1a-DIO-Flpo-WPRE-hGHpA (1×10^10^ vg/mL, Addgene). For two-photon glutamate imaging of semi-sparsely labelled L5b ET cells we injected 70 nl of AAV-fDIO-iGluSNFR-GPio (1×10^12^ vg/mL, Charite Vector Core) combined with AAV9-EpEF1a-DIO-Flpo-WPRE-hGHpA (1×10^11^ vg/mL). For two-photon Ca^2+^ imaging of L1 NDNF interneurons we injected 100 nl of AAV-hSYN-flex-Gcamp6f (1×10^12^ vg/mL, Charite Vector Core). For optogenetic activation of L1 NDNF interneurons we injected 200 nl AAV-CAG-Flex-hChR2(H134R)-mCherry (1×10^13^ vg/mL, Charite Vector Core). For experiments described in Fig 2 we targeted L5b neurons by injecting 200 nl of a retrograde virus AAVrg-EF1a-Flpo (1×10^13^ vg/mL, Addgene) in the medulla (−6.6 mm AP, 0.9 mm ML, -4.5 mm DV) and 70 nl of AAV-Ef1a-fDIO-GCaMP6s (1×10^12^ vg/mL, Addgene) in ALM. After injection, the craniotomy was sealed with a custom-made stainless-steel window barrel with a diameter of 3-mm and a depth of 0.4 mm glued to a glass coverslip (3mm). Sealing was achieved with resin cement (Relyx). Finally, the headpost was implanted using Relyx and Paladur subsequently until the complete skull was sealed. Imaging experiments began 3-4 weeks after the virus injection.

For optogenetic inhibition of the whole ALM area (Fig 1B, C, Fig S2) no cranial window was implanted. Instead, the intact skull was cleared using a layer of cyanoacrylate (Zap-A-Gap CA+, Pacer technology) to clear the bone (*42*). Headpost implantation was performed as described above.

For electrophysiological recordings in vivo we performed a craniotomy and implanted a steel headplate, a ground screw, and a 3D-printed recording chamber. After removing the skin and periosteum from the dorsal skull, the dorsal parts of the temporalis and occipitalis muscles were detached. A U-shaped steel headplate was attached to the parietal bone using resin cement (3M, RelyX). A small craniotomy was made over the cerebellum with a drill, and a steel ground screw, attached to a silver wire with a gold connector, was secured into the skull. Another 1 mm craniotomy was made over the anterolateral motor cortex (2.5 mm AP, 1.5 mm ML from bregma) using a biopsy punch (Stiefel). Silicone gel (Cambridge Neurotech, Dura-Gel) was applied to the dura to seal the craniotomy and allowed to cure for 15 minutes. The base of a recording chamber made from methacrylate compound was implanted onto the skull with dental cement (Lang Dental, Ortho-Jet), and a cap was used to seal the chamber. Animals were given a 3-hour recovery period before proceeding with the experiment.

### Behavior

One week after surgeries mice were handled and habituated. Head fixation time gradually increased. Once animals were habituated to head fixation behavioral training started. Behavior was controlled with Bpod State Machines (Sanworks) and the corresponding custom written MATLAB code. Animals were water restricted either by absence of a water bottle in their cage or by replacement of the water bottle with a 1-2% citric acid water bottle, which also resulted in increased thirst of the animals securing adecuate task involvement (*43*). Each trial in the behavioral paradigm lasted 6 seconds with 5 different trial epochs: 1.2 s initial delay, 0.7 s instruction by mildly (0.8bar) air-puffing the right or the left whisker pad, 1 s delay period in which the animal had to withhold licking, 0.1 s auditory 10kHz go-cue and finally 3 s report period where mice had to lick the corresponding licking port for a saccharine (0.1 %) water reward of 5 μl delivered after tongue contact. Licking was tracked by custom built capacitive touch detectors. Several infrared cameras (Basler) were used to track further behavior. Inter-trial-interval duration lasted 6-8 s. First animals were trained to perform the bidirectional licking task (Rule A) ignoring whether they licked during the delay period. Once animals robustly associated the instruction side with the corresponding reward port, we started punishing animals with trial abortion when licking happened before the go-cue. After animals reached about 80% of correct performance, we measured several baseline days for this task (rule A) before starting the relearning paradigm. Here animals had to switch to a unidirectional licking task where regardless of which site was instructed by the airpuff, they had to lick to the left reward port. Animals learned this within one session (day 2), which was then followed by a further session of this new task to confirm robust learning (day 3). Afterwards mice had to relearn back to rule A (A’) within one session (day 4), also followed by a confirmation session to proof relearning (day 5). The relearning paradigm lasted overall five sessions. To keep mice from stopping to perform, during relearning sessions (day 2 and 4), animals received an automatic reward on a random subsample of 10% of the relearned trials.

To quantify the correct choice performance, impulsive (lick too early) and omission (no lick) trials were discarded. However, the licking direction in impulsive trials (which happened seldomly in trained animals) was taken into account for reconstructing relearning trajectories (Fig 1 F, G) as these trials were also indicative of potential learning in previous trials.

### Two photon imaging

Imaging from behaving mice was performed with a resonant-scanning, two-photon microscope (Thorlabs) equipped with GaAsP photomultiplier tubes (Hamamatsu). GCaMP sensors were excited at 930 nm (typically 30–40 mW at the sample) for functional imaging and 860 nm for structural imaging, with a Ti:Sapphire laser (Mai Tai eHP DeepSee, Spectra-Physics) and imaged through a 16×, 0.8-numerical aperture (NA) water immersion objective (Nikon). Full-frame images (512 × 512 pixels^2^) were acquired from apical tuft dendrites of sL5b neurons expressing GCaMP6s or from NDNF interneurons expressing GCaMP6f at a depth of 50–100 µm at 30 Hz using ScanImage software (Vidrio Technologies). Two plane imaging was performed with a piezo actuator (Physik Imstrumente) with a plane distance of 60 μm resulting in a 10Hz imaging frequency for both planes. At the beginning of each imaging session a structural z-stack was acquired and saved to ensure perfect FOV alignment on the subsequent imaging session. For daily FOV alignment we used the algorithm provided by the Scanimage software. We also used this tool to track and prevent slow continuous tissue and focal plane shift during imaging sessions.

For imaging experiments measuring glutamate activity, acquisition was done at 100 Hz due to the faster dynamics of glutamate sensor iGluSNFR3 (*44*). To achieve this, we draw rectangular FOVs of interest that ensured the corresponding frame rate (normally 200×150 pixels^2^; 90 × 75 μm^2^). Sensor excitation was done at 960 nm (60 mW at sample) and different FOVs were imaged each day to prevent the glutamate sensor from bleaching. Moreover, for all mice, at least one behavioral session was recorded using a wavelength of 860 nm, which is near the isosbestic point of iGluSnFR (*32*). This resulted in fluorescence intensity being independent of glutamate concentration under this wavelength. Hence, transients visible under these experimental conditions should be solely produced by motion artefacts. We confirmed to not observe any of the behavioral activity patterns described in Fig3 under these conditions.

### Optogenetics

To perform optogenetic stimulation fiber optic cannulae (200 µm diameter, 0.39 numerical aperture, THORLABS) were used. Light intensity was controlled by a LED-Driver (ThorLabs LEDD1B) coupled to a 470nm LED (ThorLabs M470F3) and the light was calibrated to shine with a power of 3,5mW on sample. Fiber tip was placed one millimeter above ALM.

NDNF activation was performed unilaterally on the right hemisphere and light was delivered in the form of a 40 Hz sinusoidal waveform. In baseline sessions, to quantify the effect of activation of NDNF interneurons in learned behaviors (Fig 1D, E), 40% of the trials were control trials, 60% were light stimulated either during the instruction, the delay or the report epoch (20% probability each).

To compare effects of activation of NDNF interneurons on flexible learning, each mice performed the rule switching paradigm twice, once under control conditions and once under optogenetic conditions (random order). Here on day 2 (Rule A to B) and 4 (Rule B to A’) once the rule switch was activated all trials were optogenetically stimulated during the entire report epoch (3 seconds).

To test whether ALM is involved in the different tasks of our rule switching paradigm (Fig 1B, C, Fig S2), light was delivered throughout the entire duration of a given trial (6 seconds) either on one hemisphere of both hemispheres of ALM. Here, 50% of the trials were control trials while the other 50% of the trials either unilateral or bilateral stimulation of ALM (25% each) was performed.

### Chemogenetics

AAV2/8-hSyn-DIO-hM3D(Gq)-mCherry (Addgene) was injected into NDNF-Cre mice. One injection (∼150 nl) was made on the ALM at a depth of 100 µm below the pial surface. Experiments began 3 weeks after the virus injection. For systemic application of deschloroclozapine (DCZ, 0.2 μg^*^g^−1^, subcutaneously) was injected after a control session of 70 trials. After 5 minutes the effect of DCZ on behavioral performance and L5b neuron activity was tested in a subsequent set of trials (∼60 trials) doing the same task (rule A).

### Electrophysiology

Recordings were made in awake, head-fixed mice using Neuropixels 1.0 (Phase 3B, *44*) electrode arrays. The probe ground was connected to a steel skull screw, and the internal tip reference was used. Probes were lowered through Dura-Gel and the dura to their final position at a depth of 3500 µm at a speed of 4 µm/s. After insertion, probes were allowed to settle for 6 min before the recording was started. Each session included a 30-minute baseline recording followed by subcutaneous administration of DCZ (0.2 μg^*^g^−1^ in PBS), and a subsequent 30-minute recording period. During recording sessions, 700 ms periods of whisker stimulation were alternated with uniformly distributed resting periods of 5-10 seconds. Electrophysiological data were recorded with SpikeGLX (https://github.com/billkarsh/SpikeGLX). The action potential band was denoised by common median referencing and band-pass filtered with a 12th-order Butterworth filter (300 – 9,000 Hz) with CatGT (https://github.com/billkarsh/CatGT). Units were spike-sorted using Kilosort 4.0 (*46*) (https://github.com/MouseLand/Kilosort) and manually curated in Phy (*47*) (https://github.com/cortex-lab/phy) to exclude multi-unit activity and noise clusters.

To reconstruct probe trajectories, probes were labeled with fluorescent dye (Vybrant DiO, Thermo Fisher Scientific) prior to recordings by applying a drop of the dye along the probe shank and moving it up and down. After the experiment mice were perfused and the brain was extracted (*see Histology for details*). Slice images were acquired using a epifluorescence microscope at 1.25× magnification, including blue (nuclei, DAPI), green (probe track, DiO), and red (viral expression, mCherry) channels. Probe trajectories were reconstructed from the slice images by aligning them to the Allen Mouse Brain Common Coordinate Framework (*48*) using Sharp-Track (https://github.com/cortex-lab/allenCCF/tree/master). The insertion depth of each probe was further confirmed through power spectrum analysis of the LFP band (https://github.com/hanhou/code_cache/tree/master/lfpSurface). All insertions were made in the right hemisphere. The position of each recorded neuron was estimated based on its depth from the brain surface along the probe trajectory.

### Retrograde tracing

One week after Fast Blue (Polysciences) application, brain sections were prepared for ex vivo imaging under a confocal microscope. First, epifluorescence images of the entire slice were acquired as an overview. Next close-up images were acquired with a confocal microscope for further inspection.

### Histology

Mice anesthesia was induced with isoflurane. Mice were transcardially perfused with cold PBS followed by 4% PFA in 1xPBS. After perfusion, the brains were removed from the skull and postfixed in 4% PFA overnight. On the next day, the brains were washed in PBS and sectioned. Coronal sections (100-μm thick) were obtained using a vibtratome (Leica). All slices were mounted with either DAPI or FarRed DAPI mounting medium.

### Two photon imaging data analysis

Image preprocessing was done using suite2p (*49*) In short, we performed rigid and non-rigid motion correction, spatial and temporal filtering, detection of ROIs and neuropil subtraction (not performed in glutamate activity imaging).

In the case of glutamate imaging, ROI detection by suite2p was visually inspected and if needed further synaptic ROIs added or discarded. The aim was to visually detect and conserve synapses with clean and visible transients in the dataset.

For all calcium imaging experiments except those described in Fig2 and 4L, fluorescence change (ΔF/F0) was event deconvolved with the cascade algorithm (*50*). Moreover, dendritic ROIs were attributed to their corresponding dendritic tree by inspecting their morphology with an acquired structural z-stack and their activity profile similarity. For data in Fig 4 A – J, several ROIs of the same dendritic tree were averaged and analyzed together.

We defined selectivity as a measure of trial type discrimination. Selectivity was calculated as follows:

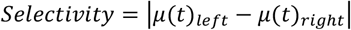

Where μ(t) is the mean activity (measured as ΔF/F0 or events/s) of right or left instructed trials. To determine whether a ROI (dendrite, synapse or interneuron) was significantly selective during a given trial epoch, we performed a Mann Whitney U test comparing their mean activity during that epoch in left versus right instructed trials. We focused on the instruction or the report epoch as selectivity peaked during those epochs. ROIs selective during the instruction and report period were labelled as ‘Both’. ROIs selective during the report period were labelled as ‘Choice’ selective.

To determine whether a ROI was modulated by transition errors we used a similar formula as for the selectivity calculation. Here only rule switching sessions (day 2 and 4) were taken into consideration. The criteria for the trial types included were as follows:

| **Transition A-B**<br>**(day 2)** | Rule A<br>(before switch) | Rule B<br>(after switch) | **Transition B-A’**<br>**(day 4)** | Rule B<br>(before switch) | Rule A’<br>(after switch) |
| --- | --- | --- | --- | --- | --- |
| *Instruction* | Right | Right | *Instruction* | Right | Right |
| *Report* | Right | Right | *Report* | Left | Left |
| *Outcome* | **Correct** | **Error** | *Outcome* | **Correct** | **Error** |

Taken together transition error modulation (T.e.m.) was calculated as:

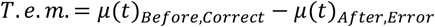

Where μ(t)_Before,Correct_ is the mean activity in correct trials before rule switch activation and μ(t)_After,Error_ is the mean activity of error trials after the rule switch activation (both under the same instruction and report conditions). In some cases, T.e.m. is reported as absolute value indicated as |T.e.m.|.

Moreover, we defined the term coding correlation as a measure of how similar two different ROIs represented the task. For this we first concatenated the mean trial activity of each ROI for two different trial types (left and right instructed) of a session. Subsequently, we measured the Pearson correlation of this vector between two different ROIs or the same ROI across different days. This can be summarized as:

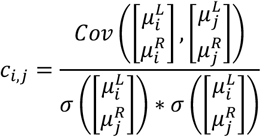

Where μ is the mean trial type activity and c is the coding correlation. For representational drift analysis *i* and *j* describe the same ROI but for different sessions i and j. In all other cases *i* and *j* represent two different ROIs.

Furthermore, we measured coactivity. Here we took the entirety of the synaptic activity traces of two synaptic ROIs and performed Pearson correlation across all time points of a session. This calculation can be described as:

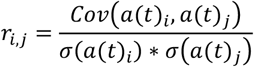

Where r_i,j_ is defined as coactivity and a(t) is the activity of i or j at any given recorded timepoint t of the imaging session.

For transient detection we first calculated noise for each ROI. We calculated noise as the standard deviation of the subtraction of the time filtered activity trace from the original raw trace.:

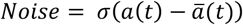

Where *a* is the activity and *ā* is the filtered activity. For *ā* activity was smoothed over 1 second. Next, transients were detected using the MATLAB function findpeaks(). The minimum peak height and minimum peak prominence were set to values at least three times greater than the calculated noise.

The start and end time points of each transient were defined as the first and last time points, respectively, where the activity trace a(t) crossed the threshold of 1*Noise before and after the detected peak.

To calculate isolated, independent calcium activity in dendritic spines, we followed the procedure described in (*26*). The activity of the parent dendritic shaft served as a reference. The ΔF/F0 of this reference was processed using a constrained deconvolution spike inference algorithm (*51*; https://github.com/epnev/constrained_foopsi_python). The algorithm was applied with an autoregressive order of 1 and a ‘fudge factor’ of 0.5.

We then fitted this reference trace to the activity traces of the spines on that same branch to construct a model that minimized the difference between the reference signal and the spine activity. The resulting model was subtracted from the raw activity trace of the spine, isolating the residual signal, which was interpreted as the independent, local activity of the dendritic spine.

The process can be described mathematically as follows:

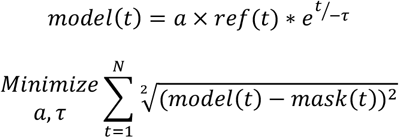

Where *a* is an amplitude constant, *τ* is the time constant for the exponential kernel and ref(t) is the foopsi-deconvolved activity trace of the parent dendrite (reference). Mask(t) corresponds to the activity trace of a given spine.

To determine the presence and the size of functional subgroups in a population of imaged NDNF interneurons we first performed PCA analysis on the ΔF/F0 activity of the imaged population of interneurons throughout a session. To detect clusters, we used a DBSCAN (Density-Based Spatial Clustering of Applications with Noise) algorithm. We specified the number of points (cells) to be determined as a dense region (functional subgroup) as two. To determine an appropriate epsilon value for DBSCAN clustering, we employed a k-distance plot analysis. This method assesses the distribution of pairwise distances between points to identify a threshold for neighbourhood density. We then computed the k-nearest neighbor distances. Following this, we sorted these distances in ascending order and plotted them to identify the elbow point—the location in the curve where the slope exhibits a sharp transition. This elbow represents the optimal epsilon value, as it marks the boundary between dense clusters and noise.

We built a linear decoder to maximize the distinction between trial types during different epochs similarly as described in (*41*). We used 50% of the trials of a given session as a training dataset and the other 50% as the test dataset. First, we defined a nx1 vector for a population of n synapses/dendrites/interneurons we calculated as follows:

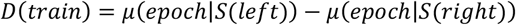

Where D(train) is a nx1 vector, and µ(epoch|S(left/right)) is the mean activity of a given ROI in left or right instructed trials during the instruction or the report epoch. We normalized the vector by its own norm. Lastly, we calculated the inner product of D(train) and the test subsample dataset (split into right and left instructed trials) to project the activity trajectories of the ROI population along D(train). Divergence between the activity trajectories of left and right instructed trials indicated a differentiation of different trial types by the population activity.

### Electrophysiological data analysis

The analysis was restricted to neurons estimated to reside in layer 5 and exhibiting an average firing rate above 0.5 Hz. To minimize the influence of drifting firing rates, comparisons between control and DCZ firing rates were made within a defined time window around the DCZ injection time (t_inj_). Baseline firing rates were calculated in the interval [t_inj_ - 5 _min_, t_inj_], while DCZ firing rates were calculated in the interval [t_inj_ + 5_min_, t_inj_ + 10_min_]. Bursts were defined as spikes preceded by an interspike interval greater than 15 ms and followed by one or more spikes with interspike intervals shorter than 15 ms. Analysis of electrophysiological data was performed in python 3.12.7.

### Statistical analysis

All statistical tests were run on MATLAB. We used nonparametric tests to avoid assumptions of normality. For paired comparisons Wilcoxon signed rank test was used, for non-paired comparisons we used Mann Whitney U tests. For multiple comparisons with more than two groups, we first run a Friedman’s ANOVA test or a one-way ANOVA test, followed by the corresponding statistical tests between single groups including Bonferroni correction. To test for linear regression of coactivity or coding correlation as a function of synaptic distance we used a Spearman correlation. All statistical comparisons are described in table S1.

To compare the effect size of DCZ manipulation on spine versus branch calcium activity, and its impact on burst versus firing rate, we calculated a ratiometric measure, where a value of 0 indicates no effect, −1-1−1 indicates a full negative effect, and 111 indicates a full positive effect. The mathematical descriptions are as follows:

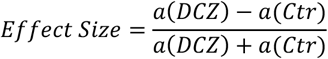

Where a(DCZ/Ctr) corresponds to the measure of activity e.g. firing rate or burst rate. To calculate the effect size in calcium imaging data shown in Fig 2, we first calculated the geometrical mean of calcium transient frequencies and amplitude to combine both aspects.

No statistical methods were used to predetermine sample sizes, but our sample sizes were like those reported in previous publications in the field.

## Supplementary Text

**Fig. S1.**
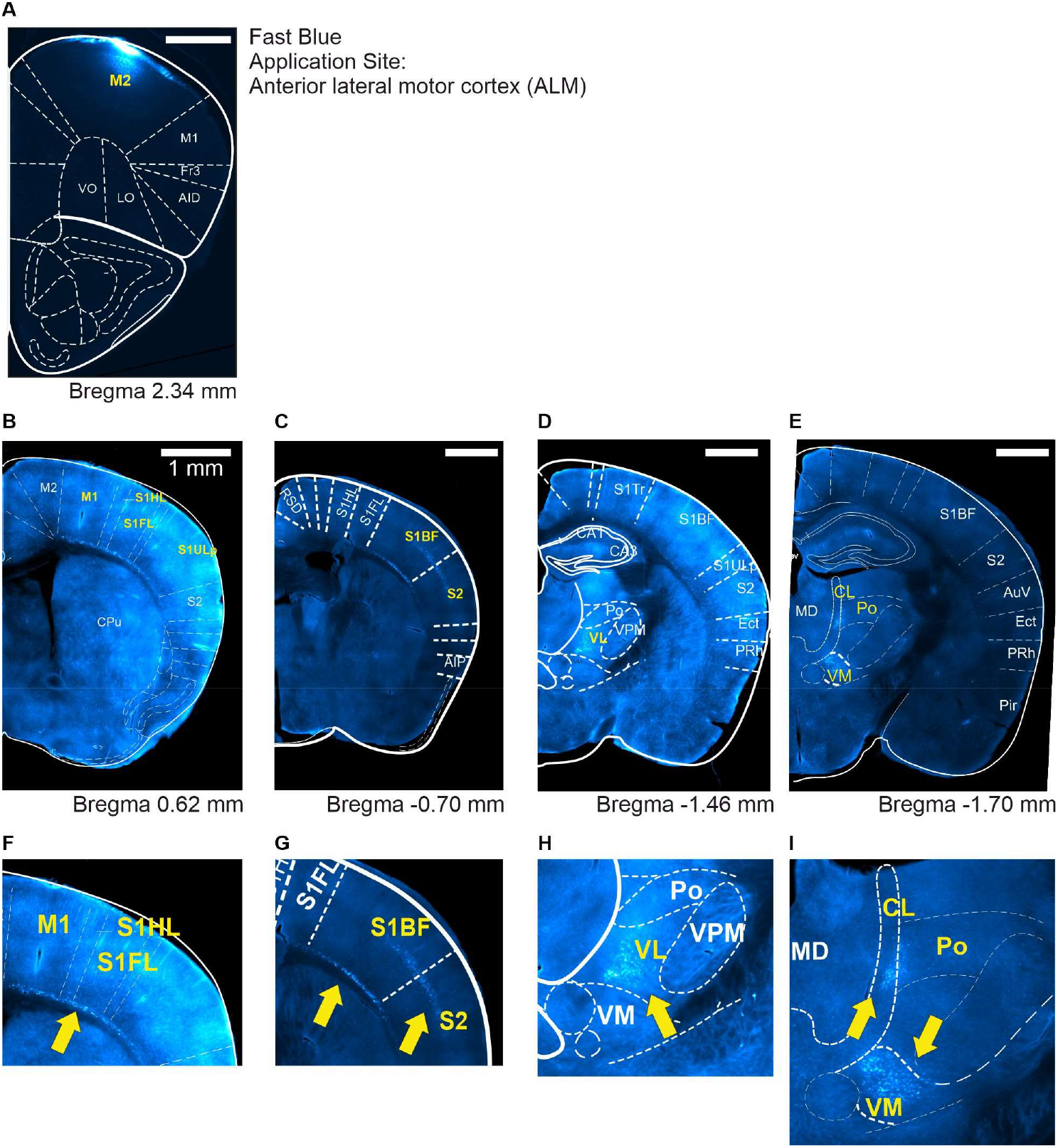
Brain regions targeting layer 1 of ALM. (**A**) Retrograde tracer fast blue was applied onto the dura on ALM. Neurons projecting to L1 will be stained with the tracer. (**B-E**) Atlas aligned coronal brain sections expressing fast blue at different positions along the anteroposterior axis. Yellow labelled brain areas indicate regions showing fast blue expression. **(F-G)** Close up view from areas expressing fast blue (see yellow arrows) aligned as in (B-E). All scale bars indicate 1 mm. Exemplary data shown from one out of three mice.

**Fig. S2.**
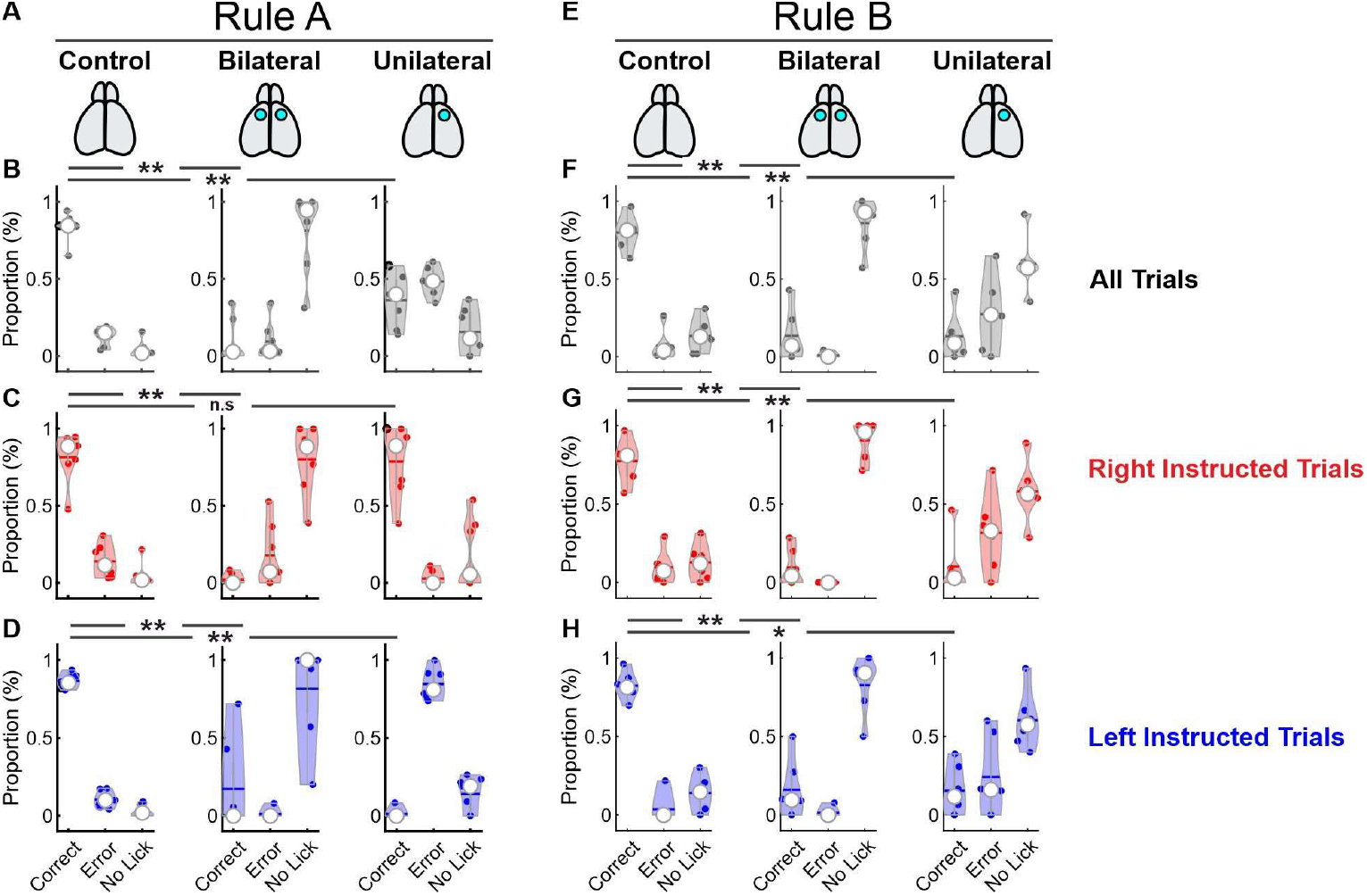
Anterolateral motor cortex enables correct task execution for both rules. **(A)** Schematic illustrating different optogenetic manipulation conditions. In Vgat-Cre × ChR2 animals, the ALM cortex was illuminated either unilaterally or bilaterally during rule A. **(B-D)** Proportion of correct, error, or omitted trials (no licking) in rule A. Left column: control trials; middle column: bilateral stimulation; right column: unilateral stimulation. **(B)** Both trial types; **(C)** right instruction trials; **(D)** left instruction trials. (One-way ANOVA followed by Wilcoxon signed rank test; see table S1; 9 sessions, 3 animals). **(E)** Schematic as in (A), but for the cue-guided simple task (rule B). **(F-H)** Proportion of correct, error, or omitted trials (no licking) in rule B. Left column: control trials; middle column: bilateral stimulation; right column: unilateral stimulation. **(F)** Both trial types; **(G)** right instruction trials; **(H)** left instruction trials. (One-way ANOVA followed by Wilcoxon signed rank test; see table S1; 8 sessions, 3 animals).

**Fig. S3.**
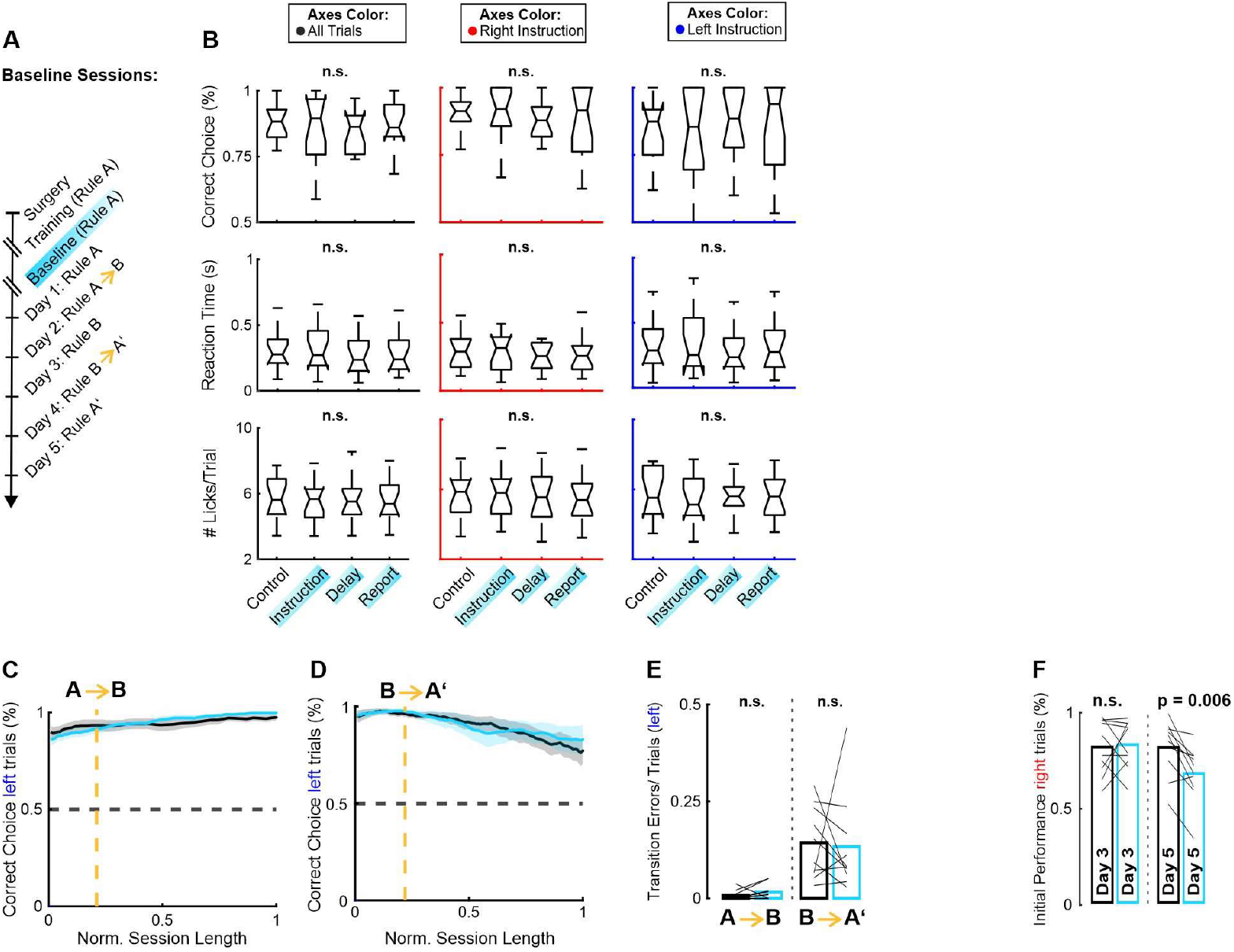
Additional quantification of optogenetic NDNF IN activation on behavior. **(A)** After animals initially learned rule A, several baseline sessions were measured while optogenetically activating NDNF interneurons in the right hemisphere of ALM at different trial epochs. **(B)** Behavioral metrics were recorded and analyzed for all trials (black axes), right instruction trials (red axes), and left instruction trials (blue axes). Metrics included correct choice percentage (top; One-way ANOVA, all p > 0.05; see table S1), reaction time (middle; One-way ANOVA, all p > 0.05; see table S1), and licks per trial (bottom; One-way ANOVA, all p > 0.05; see table S1). Data were collected across 21 sessions from 10 animals. **(C)** Left: Moving average (blocks of 20 trials) of task performance during the transition from rule A to B on left instruction trials. The orange dashed line indicates the rule change. Control condition (black) versus optogenetic manipulation (cyan). **(D)** Same analysis for the transition from rule B to A. **(E)** Transition errors per trial during left instruction trials across rule-switching sessions. Control condition (black) versus optogenetic manipulation (cyan). (Wilcoxon signed rank test: A-B, p = 0.211; B-A, p = 0.322; 10 animals). **(F)** Initial performance (correct choice; first 50 trials) on day 3 (rule B) of the relearning paradigm. Data from right instruction trials only. (Wilcoxon signed rank test: Day 3, p = 0.912; Day 5, p = 0.006; 10 animals).

**Fig. S4.**
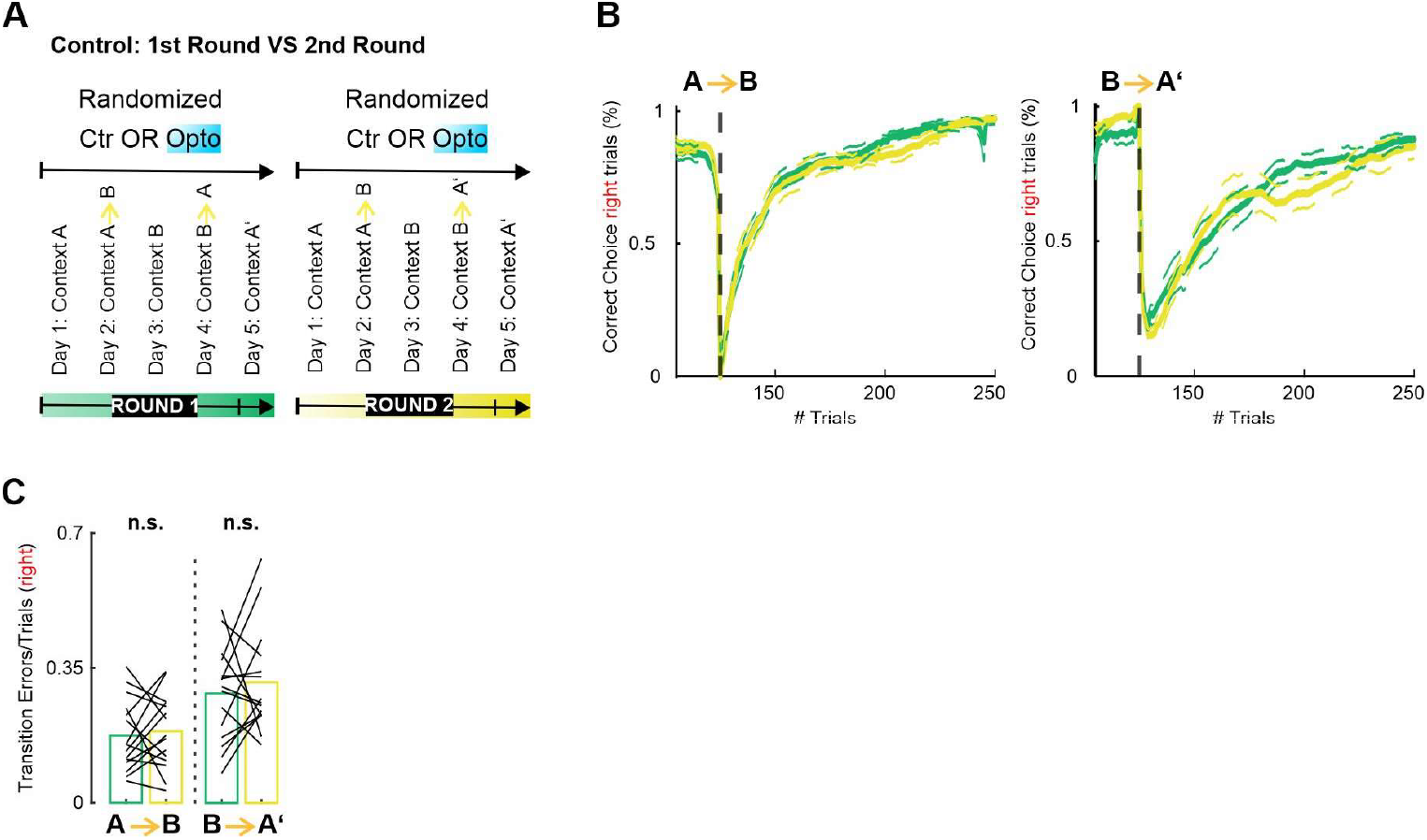
First and second relearning in the same animal iteration do not differ from each other. **(A)** Each animal completed the relearning paradigm twice: once under control conditions and once with optogenetic manipulation. The order of conditions was randomized. **(B)** Moving average of task performance (blocks of 20 trials) during transitions from Rule A to B (*left)* and B to A (*right*). Black dashed lines mark the moment of rule switch. Only right instruction trials were analyzed. The first relearning iteration is shown in green, and the second in yellow. Data are presented as mean ± SEM. **(C)** Transition error rates for right instruction trials during different rule-switching sessions. Data are shown for the first relearning iteration (green) and the second relearning iteration (yellow). Statistical analysis: A-B switch (Wilcoxon signed-rank test, p = 0.639) and B-A’ switch (Wilcoxon signed-rank test, p = 0.525). Data are from 30 relearning iterations across 15 mice.

**Fig. S5.**
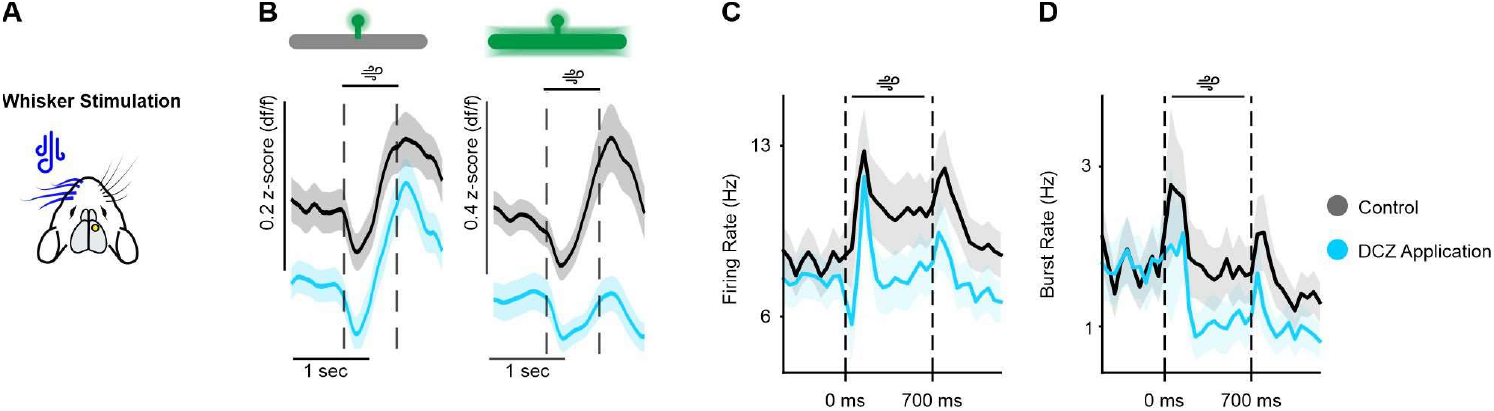
Additional quantification of the effects of chemogenetic NDNF interneuron activation on L5 neuron activity. **(A)** 2P calcium imaging and extracellular recordings were recorded while stimulating the contralateral whiskerpad. **(B)** *Left*: Average isolated spine activity under saline (black) and DCZ (cyan) application. Dashed lines indicate start of instruction period and the end of instruction period. *Right:* same for shaft events (52 branches, 122 spines; 3 mice). **(C)** Exemplary population PSTH reporting action potential firing rate during whisker stimulation under control (black) and DCZ (cyan) conditions. mean±s.e.m. **(D)** Same as (C) for burst rates. mean±s.e.m.

**Fig. S6.**
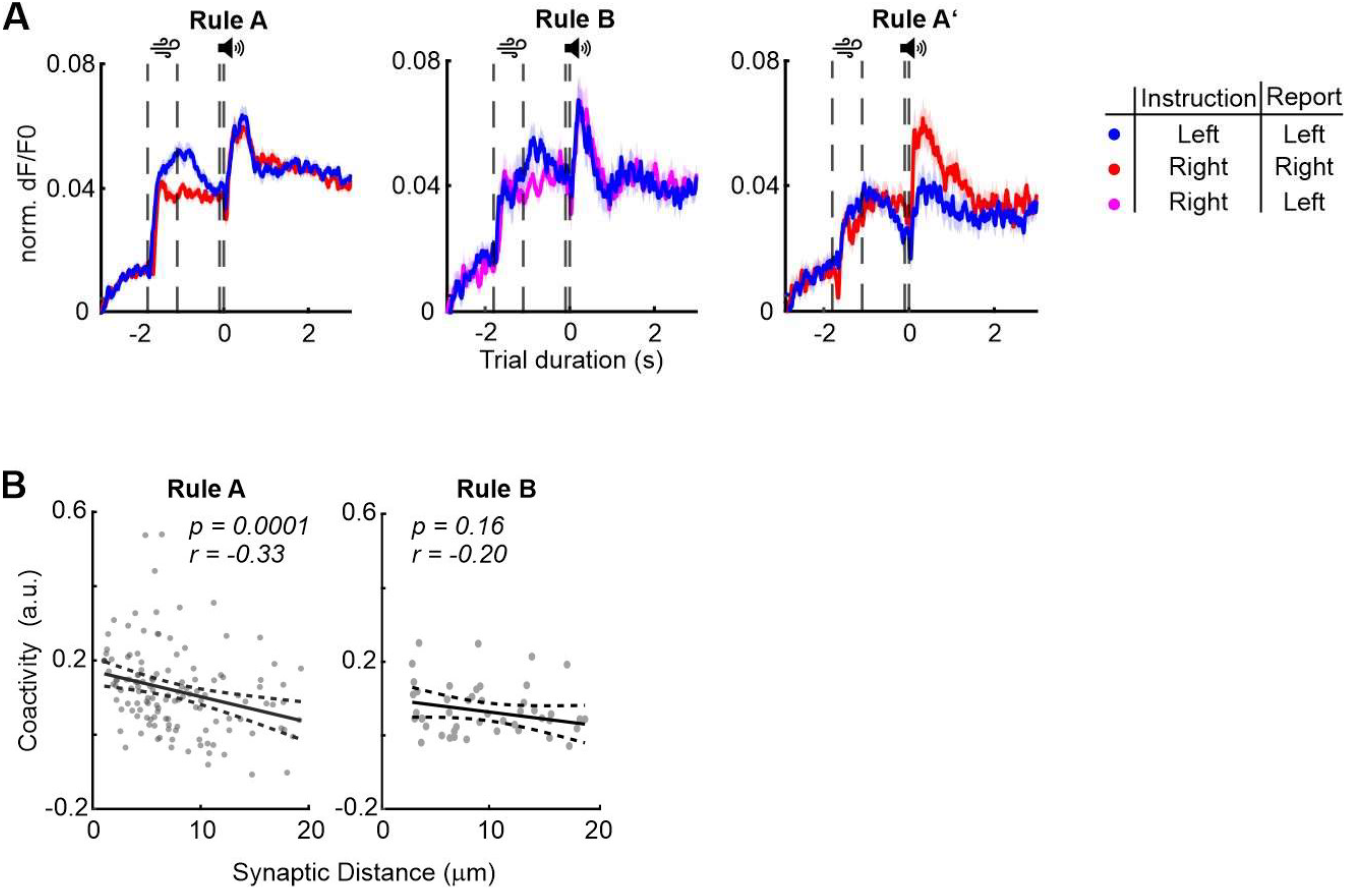
Additional analysis of synaptic glutamate imaging in apical dendrites of L5b neurons. **(A)** Average synaptic activity for different trial types across rules in the relearning paradigm. Data are shown as mean ± SEM. **(B)** Coactivity (see Methods) between synapses as a function of their distance along the dendritic branch. **Left:** Rule A (Spearman correlation, p = 0.0001; 131 synaptic pairs). **Right:** Rule B (Spearman correlation, p = 0.16; 38 synaptic pairs). Data are from 6 animals.

**Fig. S7.**
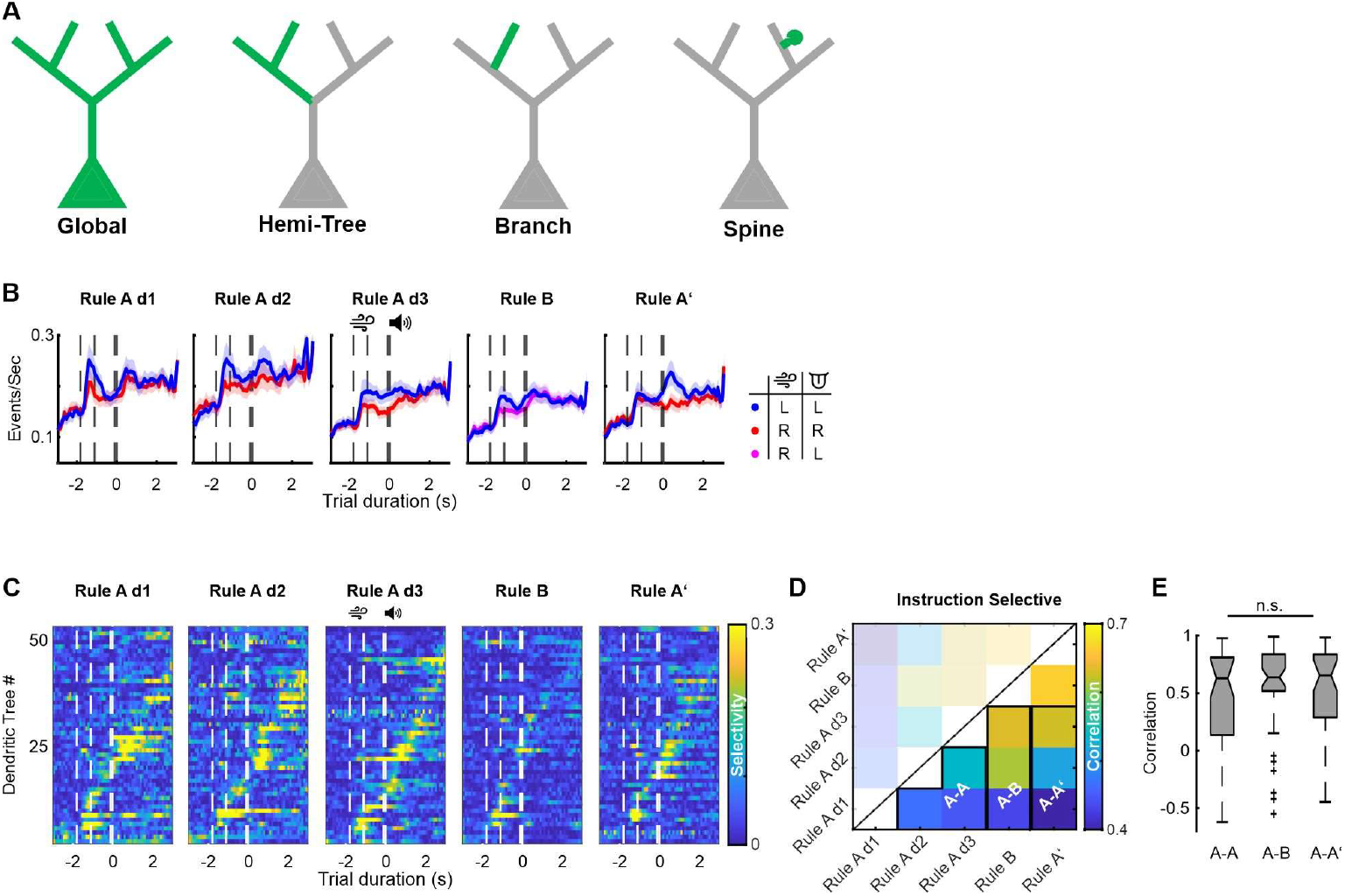
Additional analysis of calcium imaging in L5b apical tuft dendrites. **(A)** Schematic defining different transients type possibilities when performing dendritic calcium imaging. **(B)** Dendritic tree average activity reported after event deconvolution for all trial types across sessions of the relearning paradigm (mean ± SEM; 51 dendritic trees, 5 animals). **(C)** Average trial-type selectivity of all imaged apical dendritic trees, sorted daily by peak selectivity time. Dashed lines indicate the start of the instruction period, the end of the instruction period, and the go cue. **(D)** Day-to-day comparison of coding correlation (see Methods) between the same dendritic trees across the relearning paradigm. Black frames highlight comparisons between Rule A-A, Rule A-B, and Rule A-A’ sessions. Only instruction-selective neurons were analyzed. **(E)** Average coding correlation between Rule A-A, Rule A-B, and Rule A-A’ sessions for instruction-representing neurons (Friedman’s Test, p = 0.71; 14 instruction-selective dendritic trees, 25 sessions, 5 animals).

**Fig. S8.**
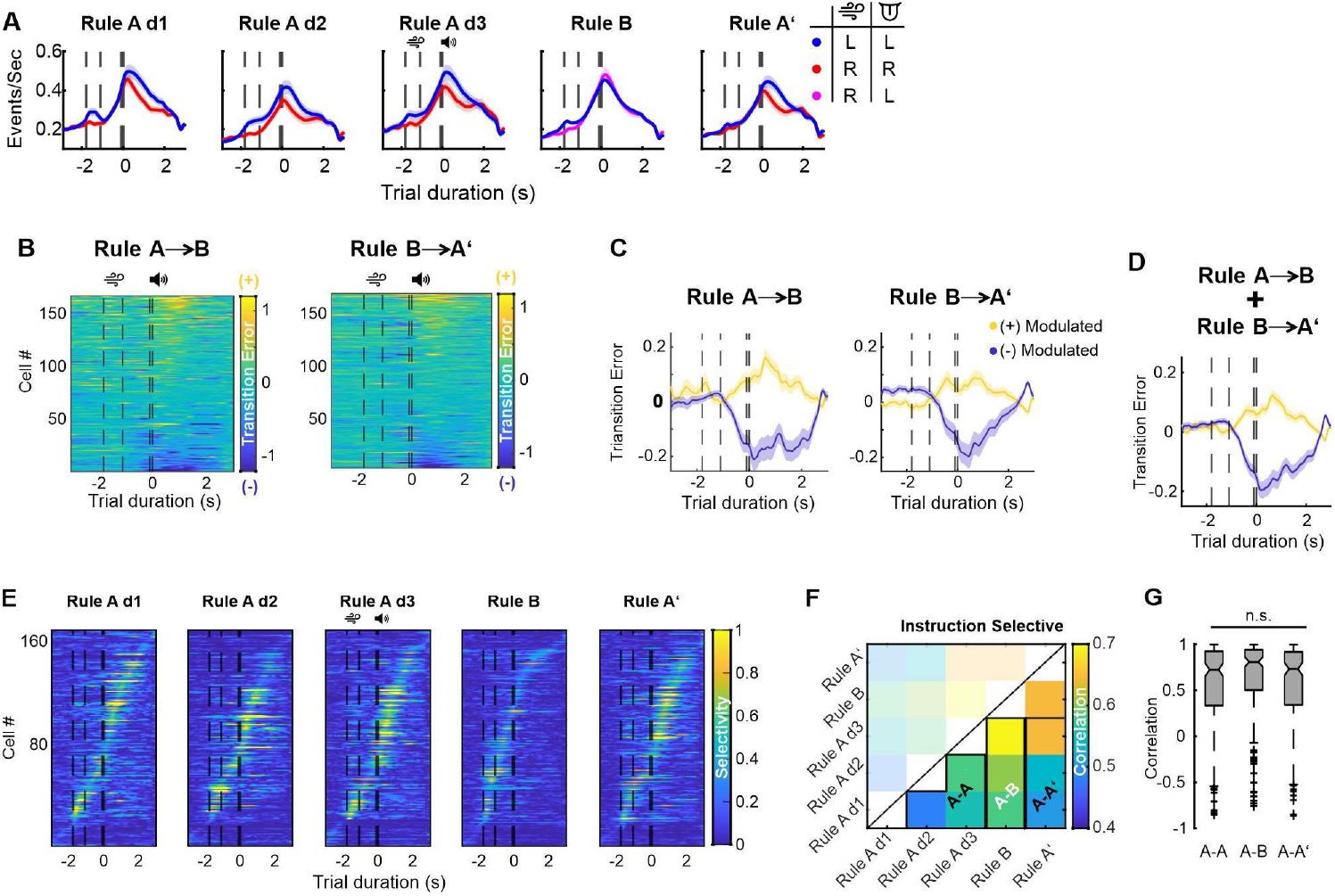
Additional analysis of NDNF interneuron imaging. **(A)** Trial averaged deconvolved calcium activity across sessions of the rule-switching paradigm. Legend indicates colors for different trial types. Mean ± SEM (170 neurons from 4 animals). **(B)** Trial averaged transition error-dependent activity for all interneurons imaged during Rule A to B (**left**) and B to A’ (**right**) transitions. Positive values indicate increased activity during transition errors, and negative values indicate decreased activity. **(C)** Average error modulation, separated into positively and negatively modulated neurons, for Rule A to B (**left**) and B to A’ (**right**) transitions. **(D)** Average error modulation for positively and negatively modulated neurons, merging data from both rule switches. **(E)** Average trial-type selectivity of all imaged NDNF interneurons, sorted daily by peak selectivity time. Dotted lines indicate the start of the instruction period, the end of the instruction period, and the go cue. **(F)** Day-to-day comparison of coding correlation (see Methods) between the same interneurons across relearning. Black frames highlight comparisons between Rule A-A, Rule A-B, and Rule A-A’ sessions. Only instruction-representing neurons were analyzed. **(G)** Average coding correlation between Rule A-A, Rule A-B, and Rule A-A’ sessions for instruction-representing neurons (Friedman’s Test, p = 0.187; 57 instruction-selective neurons, 30 sessions, 4 animals).

**Fig. S9.**
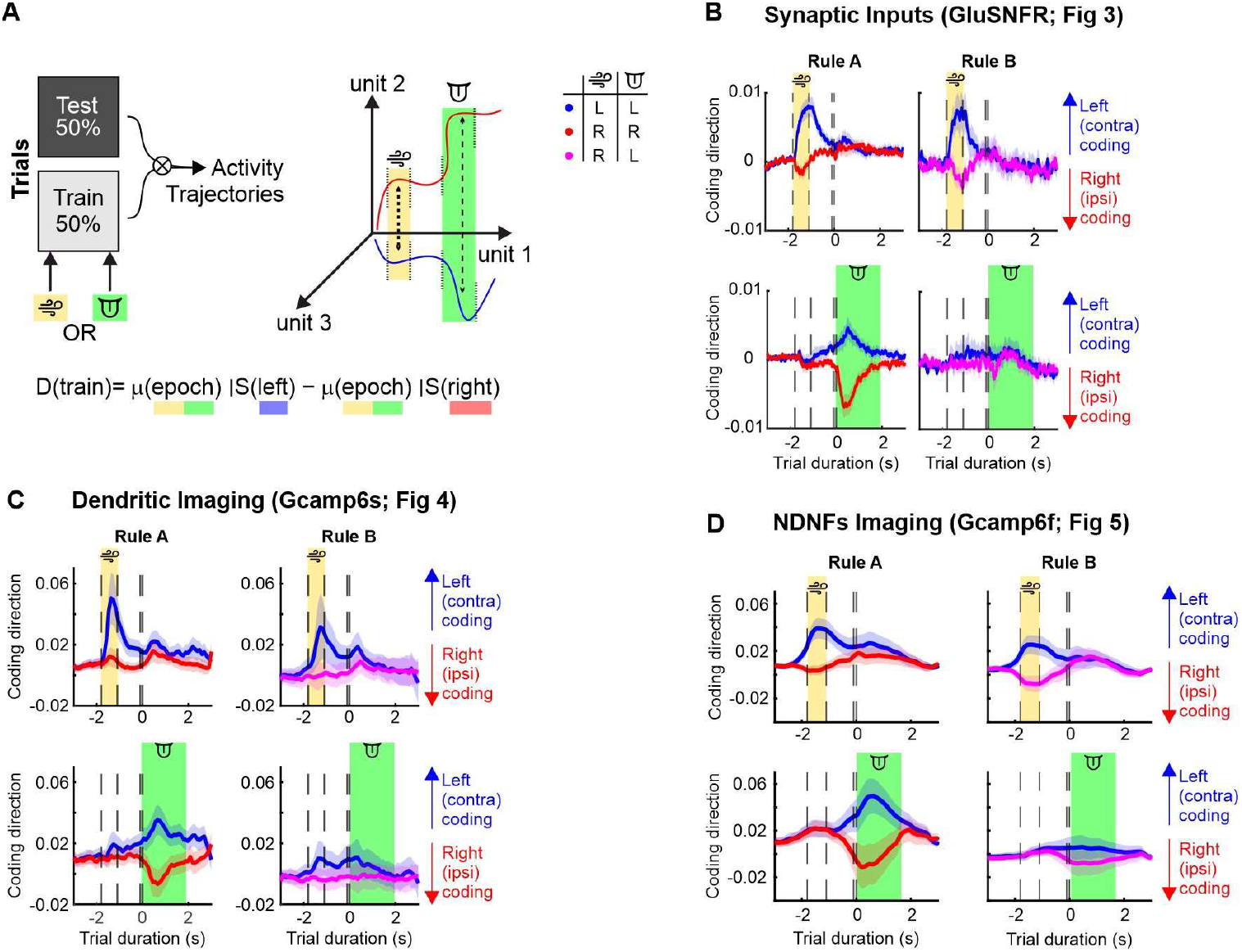
Linear decoder analysis for different circuit components. **(A)** Schematic for the linear decoder analysis. To analyze lateralization (contra- or ipsilateral) of task representation we builta linear decoder. Fifty percent of trials were used for training. The training subset was extracted from either the instruction or report epoch (shaded areas). Activity trajectories for left and right trials (test data) were projected along the time axis based on calculated trial-type discrimination D(train) during the selected epoch. **(B)** Activity trajectories for different trial types, as illustrated in the schematic. **Left column:** Rule A. **Right column:** Rule B. **Upper row:** Linear decoder trained with data from the instruction period. **Lower row:** Linear decoder trained based on data from the report period (527 synapses from all sessions, 6 animals). **(C)** Same as (B), but for dendritic calcium imaging. (51 dendritic trees, 5 animals). **(D)** Same as (C), but for NDNF interneurons (170 neurons, 4 animals).

### Interpreting the linear decoder (Fig S9)

To further decipher how the different circuit components investigated in this paper (Fig 3, 4, 5) lateralized the representation of different task epochs (instruction or response epoch), we built a linear decoder (see Methods, Fig S9, *41*). Our decoder confirmed that all of them discriminate trial types during the instruction period throughout rules. Here, contralateral airpuffs (left instruction) are represented stronger than ipsilateral ones as left (blue) activity trajectories deflect stronger from 0 than right instruction (red) activity trajectories (Fig S9). In contrast, all circuit components discriminate trial types during the report epoch only in rule A but not B. Concretely, in rule A contra- and ipsilateral choices are represented with similar strength as both left(blue) and right (red) activity trajectories deflect from 0 with a similar magnitude (Fig S9).

### Detailed description of results on ALM inhibition during behavior (Fig S2)

We inhibited ALM throughout the trial duration (6 seconds) uni- and bilaterally during either rule A or rule B (Fig S2 A, E). We subdivided our analysis into three groups: performance quantification including all trials (Fig S2 B, F), right instructed trials (Fig S2 C, G) or left instructed trials (Fig S2 D, H). Under control conditions mice showed expert performance under both rules (Fig B – D, F – G; first column). Bilateral inhibition of ALM prevented mice from licking, robustly increasing the omitted trials, and thus drastically reducing performance (Fig B – D, F – G; second column). Moreover, in rule A, right hemisphere unilateral inhibition did not decrease performance on right instructed trials but markedly induced wrong licks (right licks) on left instructed trials (Fig S2 C, D; third column). On the other hand, in rule B, right hemisphere unilateral inhibition mainly prevented the animals from licking on both trial types (right airpuff – lick left; left airpuff – lick left). Thus unilateral stimulation induces a significant drop in performance for both rule A and B but generates different types of error (rule A, wrong side lick; rule B, no lick)

**Table S1.**
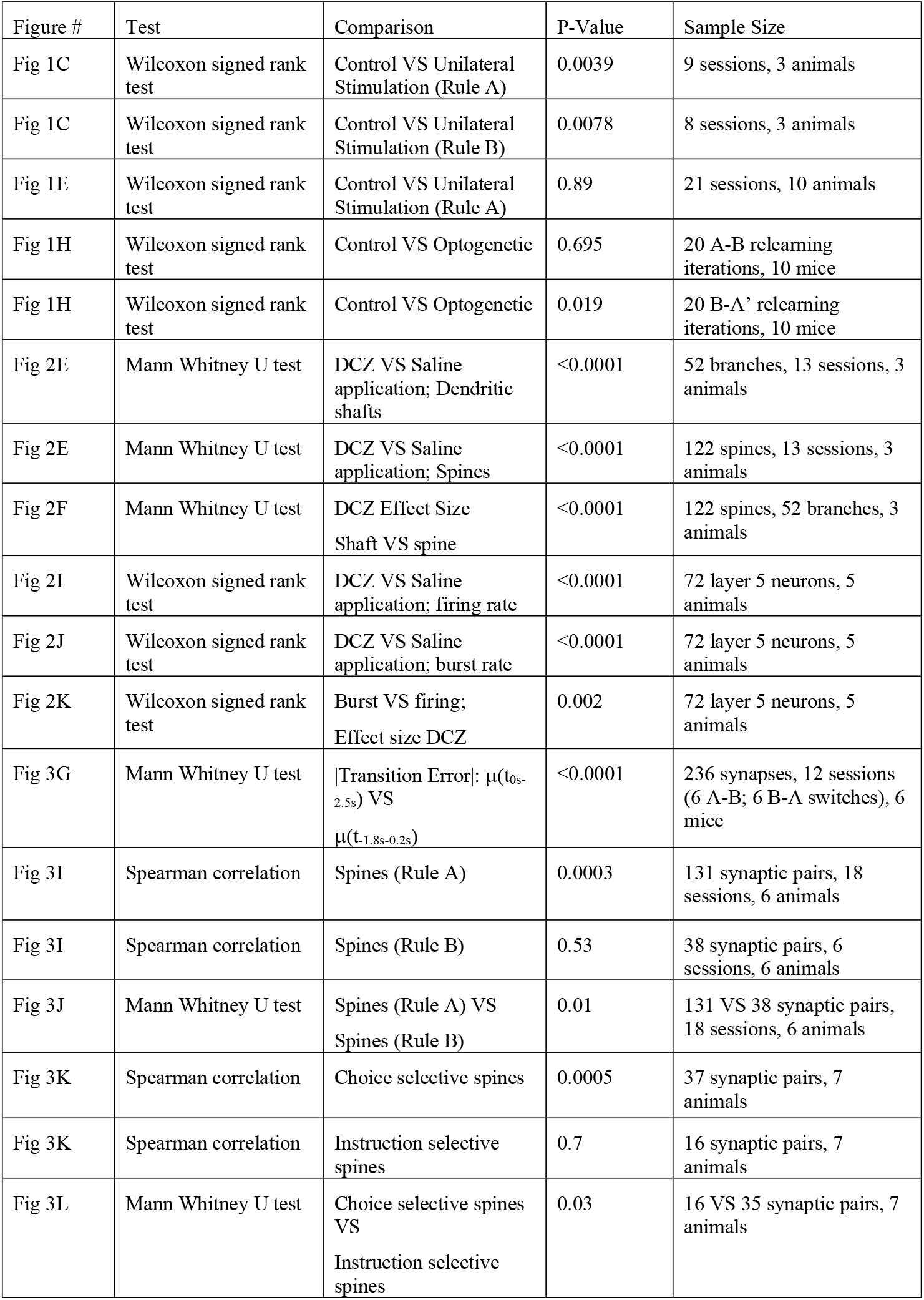
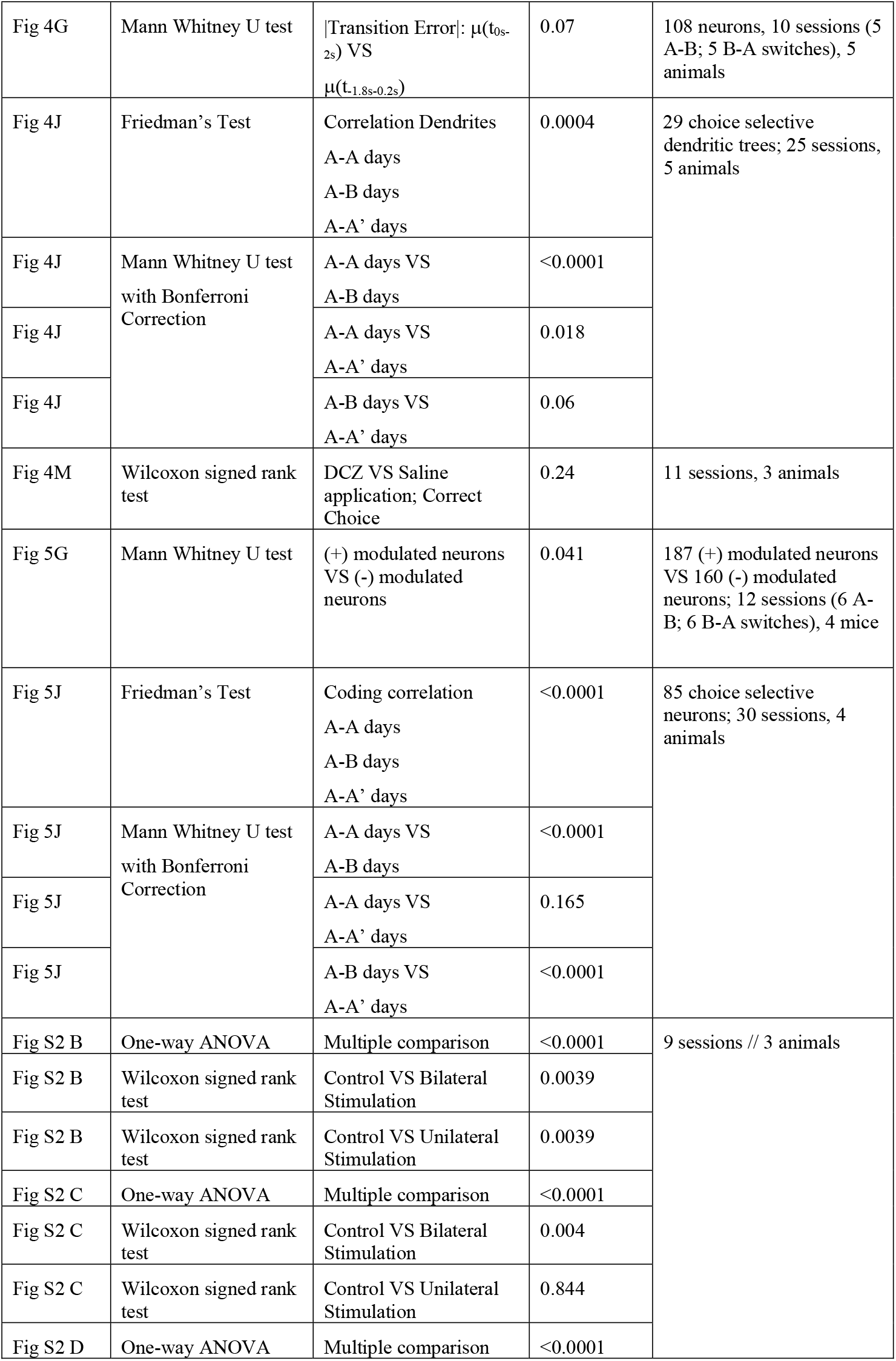
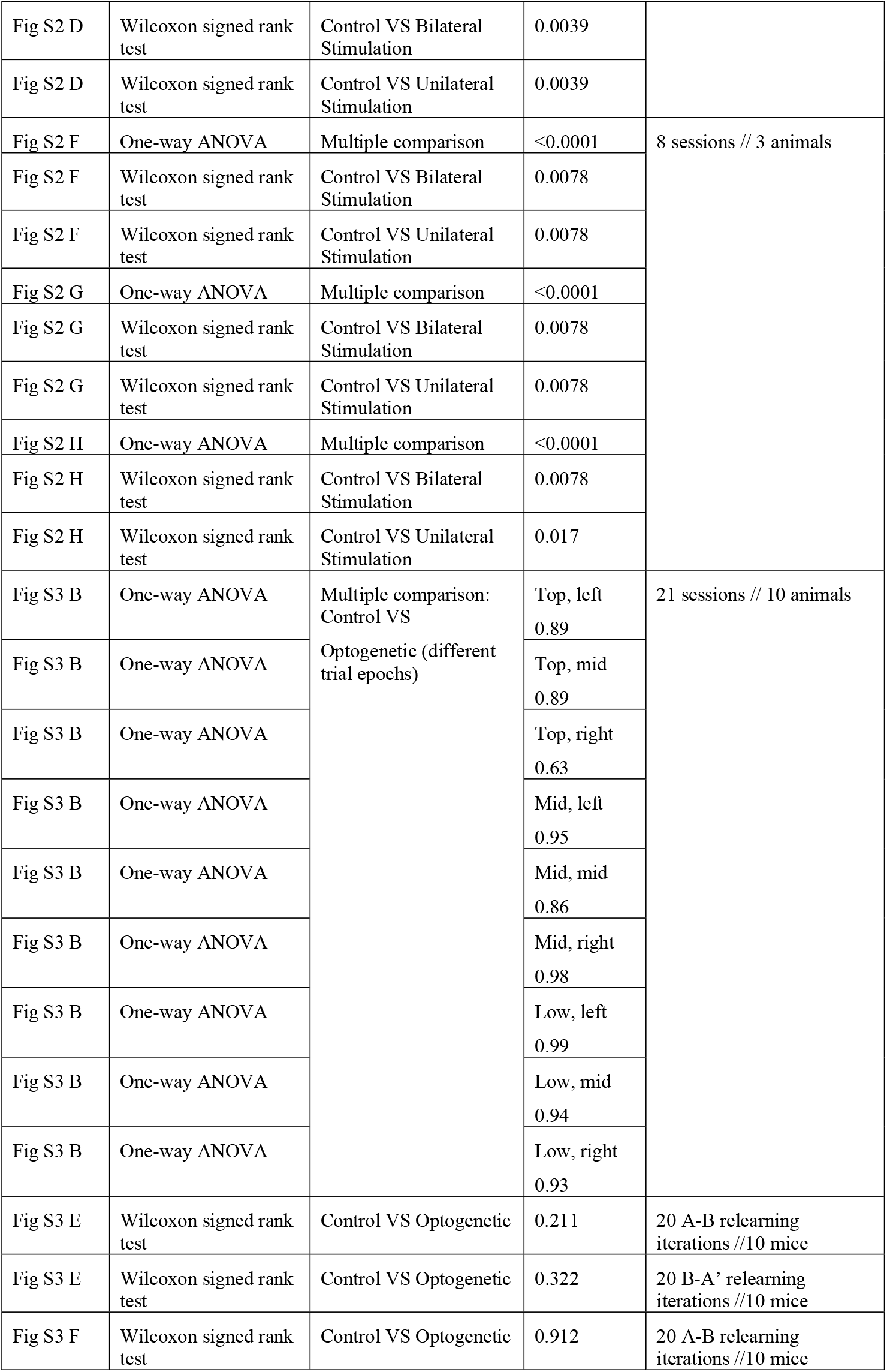
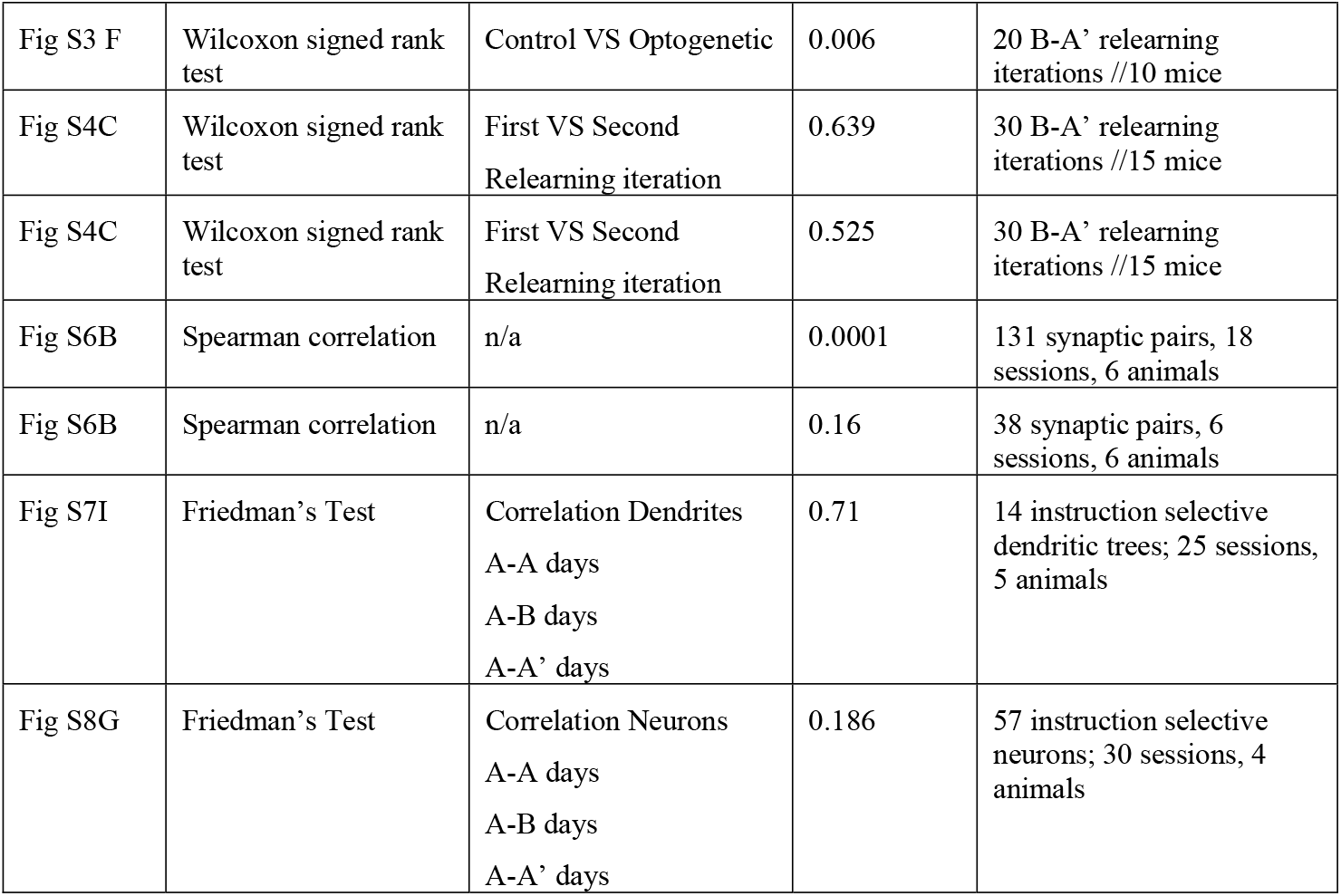
P-values and sample size for all statistical comparisons.

